# RNA degradation modulates unique aging-related gene expression in naked mole-rats

**DOI:** 10.1101/2025.11.28.691058

**Authors:** Ryuma Matsubara, Jinyu Wu, Atsuko Nakanishi Ozeki, Yoshimi Kawamura, Michiaki Hamada, Kyoko Miura, Nobuyoshi Akimitsu

## Abstract

Gene expression is co-regulated by the rates of RNA synthesis and degradation, and recent evidence has linked their imbalance to the onset of aging. The naked mole-rat (NMR) is a small rodent with an exceptionally long lifespan and markedly delayed aging, yet little is known about its RNA synthesis and degradation characteristics compared to other rodents that age more rapidly. Here, we investigate RNA synthesis and degradation in NMR and mouse skin fibroblasts by monitoring incorporation of a uridine analog, 4-thiouridine. Cross-species analysis showed that the NMR cells have higher overall rates of RNA synthesis and degradation. It further revealed higher RNA degradation rates in aging-related pathways, notably Mtorc1 signaling, likely contributing to reducing the overall expression. Although known aging-associated genes, including *Xrcc5, Nudt1, Fen1*, and *Aptx*, were expressed at similar levels in NMR and mouse fibroblasts, the RNA turnover rates were largely altered. To uncover the underlying mechanism enabling differential control of RNA kinetics, we analyzed the transcript feature importance by machine learning and identified key features governing RNA degradation both common and unique in NMR and mouse fibroblasts. Our data highlight a potential role of RNA synthesis and degradation as hidden layers of gene regulation in NMR.

## Introduction

Aging is a fundamental, steady reduction of physiological function associated with age (Stewart et al., 2005). Aging is multifaceted and its drivers scatter across virtually all aspects of cellular function (López-Otin et al., 2023), but it is well accepted that changes in gene expression are the major drivers of aging. Generally, expression changes are interpreted as the outcome of altered RNA synthesis rates, and so transcriptional regulators such as DNA methylation, histone density or distribution (heterochromatin stability), and histone modifications, have been extensively studied in aging (Wu et al., 2024). However, it is increasingly clear that gene expression is not governed by just the synthesis rate, but by the balanced acts of transcription (RNA synthesis) and clearance (RNA degradation) (Kawata et al., 2020; Timmers & Tora, 2018; Unruh et al., 2024). RNA is not inherently unstable; rather, its degradation is under tight control so a specific RNA receives selective degradation (Dowdle & Lykke-Andersen, 2025; Liu et al., 2021; Rambout & Maquat, 2024; Schoenberg & Maquat, 2012; Yamada et al., 2020). Consequently, the temporal control of RNA amount via regulation of synthesis and degradation, termed RNA kinetics, serves as a key layer of transcriptome regulation (Kawata et al., 2020).

As such, RNA kinetics is likely associated with the aging process. In fact, genes involved in RNA degradation are downregulated in senescent cells (Koh et al., 2025; Mullani et al., 2021; Sugawara et al., 2022) and artificial downregulation of RNA degradation factors causes pluripotent cells to pre-senescence (Han et al., 2024). Moreover, experimentally boosting the RNA surveillance machinery elongates the lifespan of nematodes, further highlighting the importance of RNA degradation for longevity (Son et al., 2017). The other pillar of RNA kinetics—RNA synthesis—is also widely recognized as a cause and consequence of senescence as mentioned above, and reviewed in (Wu et al., 2024).

Despite mounting evidence, the current knowledge of RNA kinetics in aging is scarce. Importantly, many studies were performed on laboratory animals (*e.g.*, mouse; *Mus musculus*) which are inherently short-lived and may have lost the mechanisms to sustain organismal longevity. In this aspect, investigating RNA kinetics in long-lived individuals together with their short-lived counterparts is a compelling strategy to identify meaningful changes. However, although the study of centenarians (humans living more than 100 years) has given us indispensable information (Kaeser et al., 2021; Santos-Pujol et al., 2025; Sato et al., 2021; Ying et al., 2024), such studies suffer from the inherent heterogeneity in the population, difficulties at setting short-lifespan control groups, as well as a lack of experimental setups for hypothesis-testing. As an alternative, cross-species studies between long-and short-lifespan mammals have the potential to identify and validate RNA kinetics as an aging factor.

The naked mole-rat (NMR; *Heterocephalus glaber*) is a subterranean rodent native to Africa (Fig. 1A). NMR is known for its exceptionally long lifespan, with a maximum lifespan of over 40 years in captivity and over 17 years in the wild (Buffenstein & Jarvis, 2002; Buffenstein, 2008; Ruby et al., 2024). NMR shows a distinctive aging pattern with no detectable age-related alterations in basal metabolic rate, gastrointestinal absorption efficiency, body composition, or bone density up to 20 years or more (Buffenstein & Jarvis, 2002; Buffenstein, 2008; Can et al., 2022; Ruby et al., 2018). NMR also shows a remarkable resistance to both sporadic and experimentally-induced oncogenesis (Buffenstein, 2008; Delaney et al., 2013; Oka et al., 2022).

**Figure 1.**
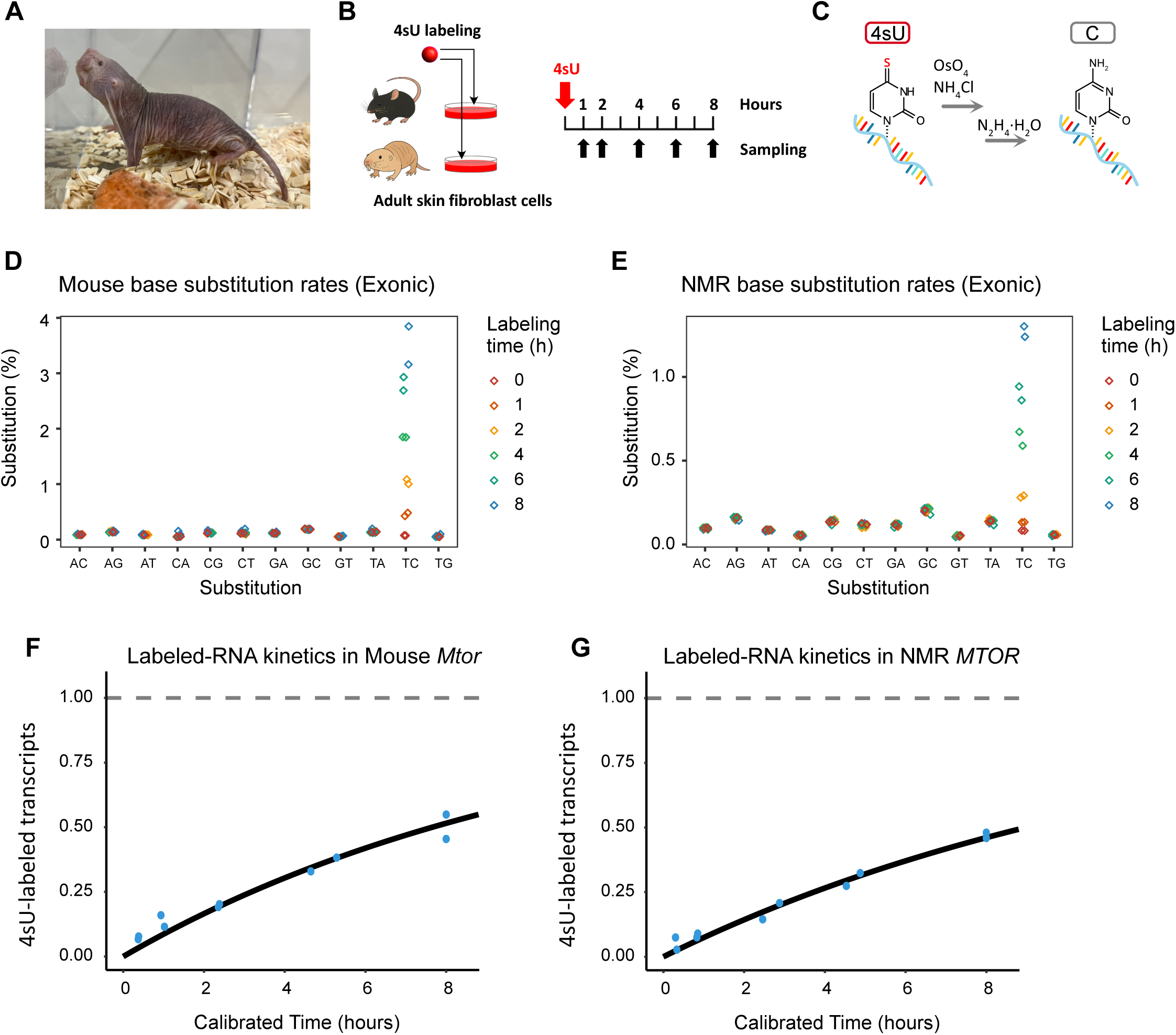
RNA metabolic labeling of naked mole-rat. (A) A naked mole-rat (NMR) maintained in our facility. (B) The experimental setup of RNA metabolic labeling. Adult skin fibroblasts of NMR and mouse were incubated in media supplemented with 100 µM 4-Thiouridine (4sU), and the RNA harvested at the indicated time points. The illustration of animals was generated by Microsoft Copilot. (C) Schematic of the TUC-seq DUAL reaction (Gasser et al., 2020). The incorporated 4sU is converted to the natural nucleobase, cytosine. The RNA samples were subsequently sequenced to identify the signature U-C substitution. OsO_4_, Osmium tetroxide; NH_4_Cl, Ammonium chloride; N_2_H_4_·H_2_O, Hydrazine monohydrate. (D, E) Substitution rates of bases in all exon-mapped reads in mouse (D) and NMR (E). Note the incremental TC substitution in both species. (F, G) Kinetic curves of mouse *Mtor* (F) and NMR *MTOR* (G) transcripts shown as examples. The rates of 4sU-labeled transcripts were calculated by GRAND-SLAM (Jürges et al., 2018) and the labeling duration recalibrated by *grandR* (Rummel et al., 2023).

Being of a similar size and classified in the order Rodentia alongside mouse, it makes an excellent model to investigate mechanisms underlying longevity and slow aging. In this study, we measured and compared the RNA kinetics in skin fibroblasts of a short-lived (mouse) and a long-lived (NMR) mammal by time-series RNA metabolic labeling experiments.

## Results

### Estimation of RNA kinetics in NMR and mouse

To measure RNA kinetics, we labeled primary skin fibroblasts of NMR and mouse (C57BL/6) with a uridine analog 4-thiouridine (4sU). Within 2 to 5 passages post-isolation, the cells were progressively labeled with 4sU, followed by sampling at the indicated time points (Fig. 1B). Total RNA was purified and subjected to TUC-Seq DUAL reaction (Gasser et al., 2020) to chemically convert the incorporated 4sU to the natural nucleobase, cytosine (Fig. 1C). The RNA was subsequently sequenced, and the reads of 4sU-containing RNA were identified by detecting the U to C mutation. The estimated rates of 4sU incorporation were up to 1.2% for NMR and 4% for mouse (Fig 1D-E), indicating sufficient 4sU labeling in both species. The RNA kinetics were then calculated by fitting the rates of newly synthesized RNA to a pre-established kinetics model with the R package *grandR* (Rummel et al., 2023). Examples of fitting are shown for NMR and mouse *Mtor* genes (Fig. 1F-G). The finalized RNA kinetics parameters including the synthesis constant (*ks*) and degradation constant (*kd*) were used for downstream analyses.

### Selecting steady-state genes for reliable kinetics estimation

The RNA kinetics estimation bases on the assumption that the synthesis and degradation rates are in equilibrium, and expression levels remain constant across the 4sU-labeling time course (steady-state; Fig. 2A). Although *grandR* is benchmarked for both steady-state and non-steady-state genes, the performance significantly suffers if used for non-steady-state genes (Rummel et al., 2023). A compelling approach is to perform differential expression (DE) analysis and filter out the DE genes; however, genes that exhibit large deviation in expression may escape detection by DE analysis and be retained in dataset (Fig. 2B, right). Therefore, to ascertain the quality of the estimation, we devised a method to detect and exclude non-steady-state genes by using 3 filters: (i) DE analysis for overall gene expression; (ii) the maximum value of replicate differences within each time point; and (iii) the maximum of the fold-changes between any combinations of the time points. The detailed explanation for the filters can be found in Supplemental Fig. 2. Genes failing any of the 3 filters were classified as non-steady (Fig. 2C). The thresholds of filters were optimized by receiver operating characteristic (ROC) analysis and grid search using the manually-curated ground truth set (Supplemental Fig. 1, also see methods). The finalized parameters (DESeq2 p.adj ≥ 0.05, difference within replicates ≤ 0.25, fold-change ≤ 1.45) achieved a Youden index (Youden, 1950) of 0.90, F1-score of 0.94, precision of 0.95, and recall of 0.95, demonstrating robust classification accuracy with high sensitivity and specificity for identification of steady-state genes. We named this filtration method as 3-layer test. After applying this test, the selected genes showed steady-state gene expression patterns, validating our method (Fig. 2D). A total of 7,281 genes (NMR) and 3,509 genes (mouse) passed both the 3-layer test and *grandR* kinetics estimation, and were used for downstream analyses (Table 1). For species comparison, 2,124 genes were recovered as common subsets between NMR and mouse genes. The kinetic parameters of all calculable RNAs are available in Supplemental Table 1.

**Figure 2.**
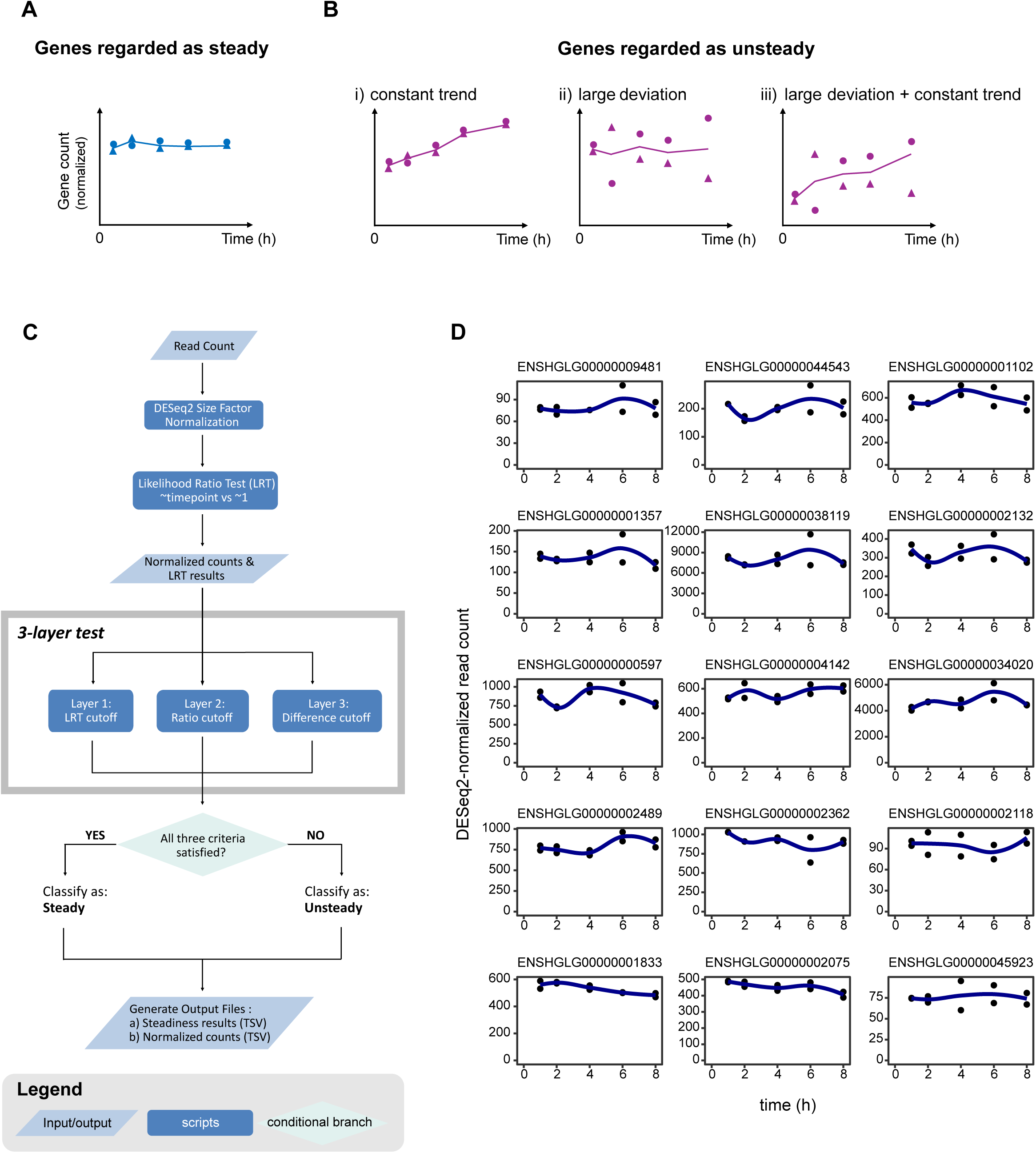
The 3-layer test to filter out non-steady-state genes. (A, B) Schematic drawings of steady (A) and non-steady (B) genes. Non-steady-state genes were defined as i) genes with expression changes; ii) with large deviation in expression; or iii) with both expression changes and large deviation. (C) A flow chart of the 3-layer test. (D) Examples of steady-state genes and their expression. Fifteen genes were randomly chosen and their DESeq2-normalized read counts were plotted through the time series. The blue lines denote non-linear regression by locally estimated scatterplot smoothing (LOESS).

**Table 1.**
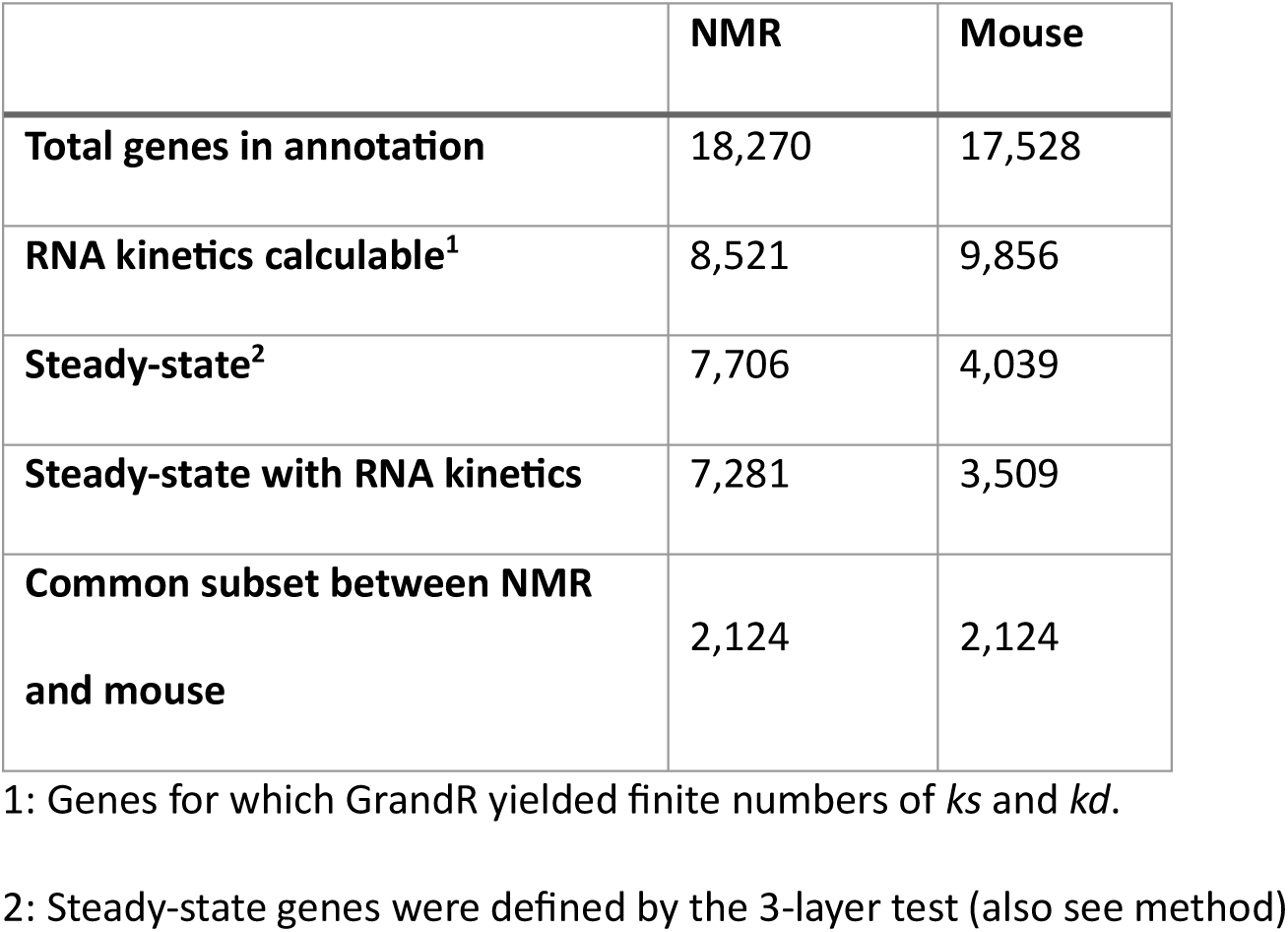
Number of selected genes in each species.

### RNA synthesis and degradation rates correlate with gene functions in NMR

In human cells, stress response genes are known to have rapid turnover rates to quickly respond to stimuli (Kawata et al., 2020). To assess the correlation between RNA turnover rates and gene functions, 7,281 genes from NMR, in steady-state and with available RNA kinetics, were divided into 9 categories according to the 1^st^ and 3^rd^ quantiles of *ks* and *kd* (Fig. 3A). Gene ontology term enrichment analysis (GO analysis) of each category revealed that genes requiring rapid response, such as DNA/transcription factor binding, show faster synthesis and degradation rates which likely ensure a swift reaction to environmental cues (Fig. 3A category C; Fig. 3B; supplemental table 2). Conversely, house-keeping genes (*i.e.*, metabolic processes) were enriched in the category of slow synthesis and degradation (Fig. 3A category G; Fig. 3B). Ribosomal structural constituents showed fast synthesis, but slow degradation (Fig. 3A category I; Fig. 3B). Faster synthesis and slow degradation result in high expression level which likely supports major cellular metabolism such as ribosome biogenesis. Overall, the RNA turnover rates correlate well with the characteristics of gene functions.

**Figure 3.**
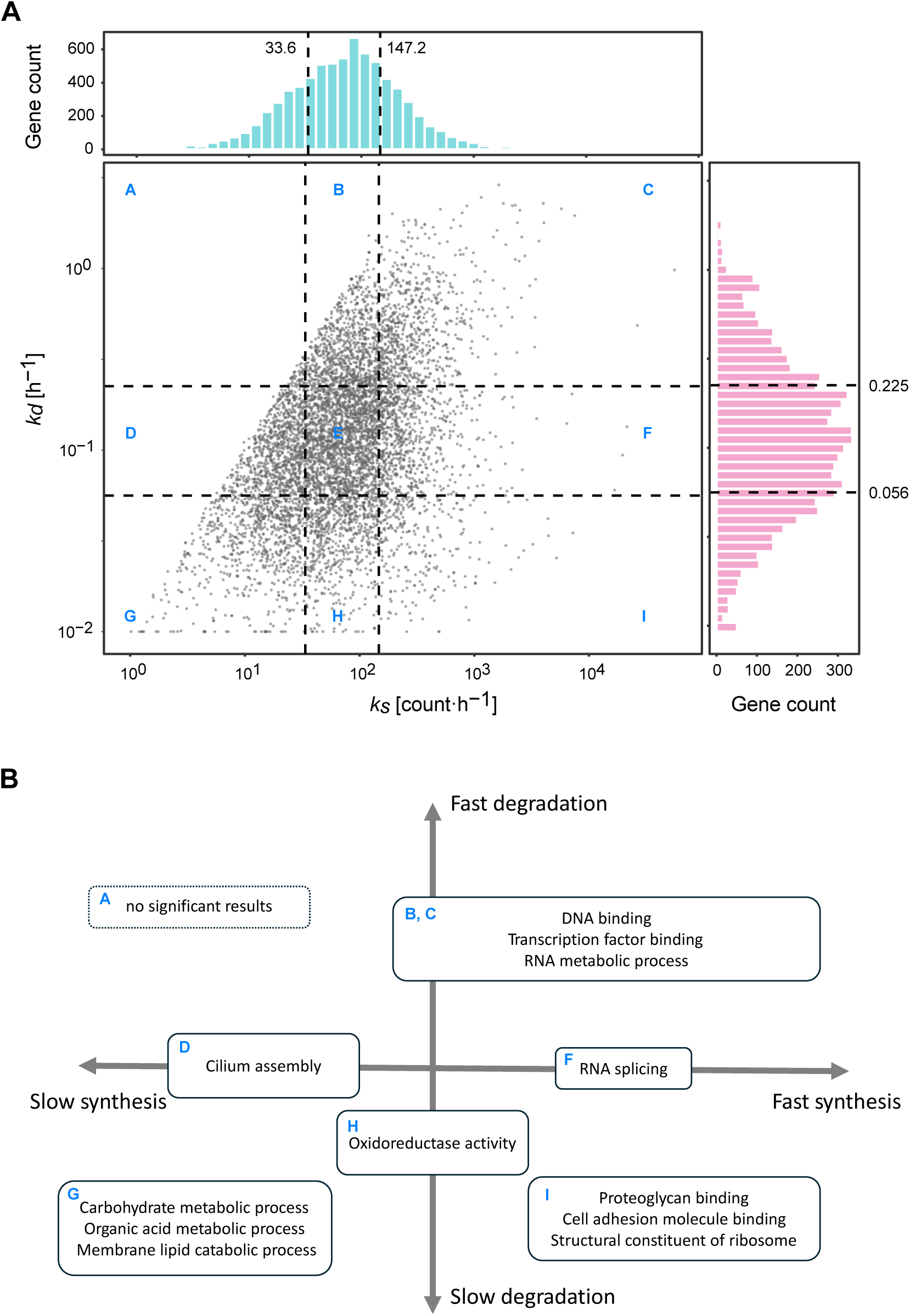
RNA synthesis (*ks*) and degradation (*kd*) rates correlate with gene functions in NMR. (A) Distribution of *ks* and *kd* in the NMR transcriptome. Steady-state genes with available RNA kinetics (7,281 genes) were analyzed. The *ks* and *kd* values were binned by the 1^st^ and 3^rd^ quantiles, and genes were categorized in 9 subgroups from A to I. (B) Representative GO-terms of genes in each category defined in (A). Pathways with adjusted *p*-value < 0.05 were considered significant. See also Supplemental Table 2 for a list of GO terms in each subgroup.

### NMR exhibits higher overall and pathway-specific RNA turnover than mouse

To understand NMR-specific gene regulation in terms of RNA turnover, we next investigated interspecies differences in RNA synthesis and degradation between NMR and mouse.

Unexpectedly, the distribution of *ks* and *kd* revealed higher overall turnover rates in NMR than that in mouse (Fig. 4A-B). This is surprising given that the protein turnover in NMR and other long-lived species is generally slower (Swovick et al., 2021).

**Figure 4.**
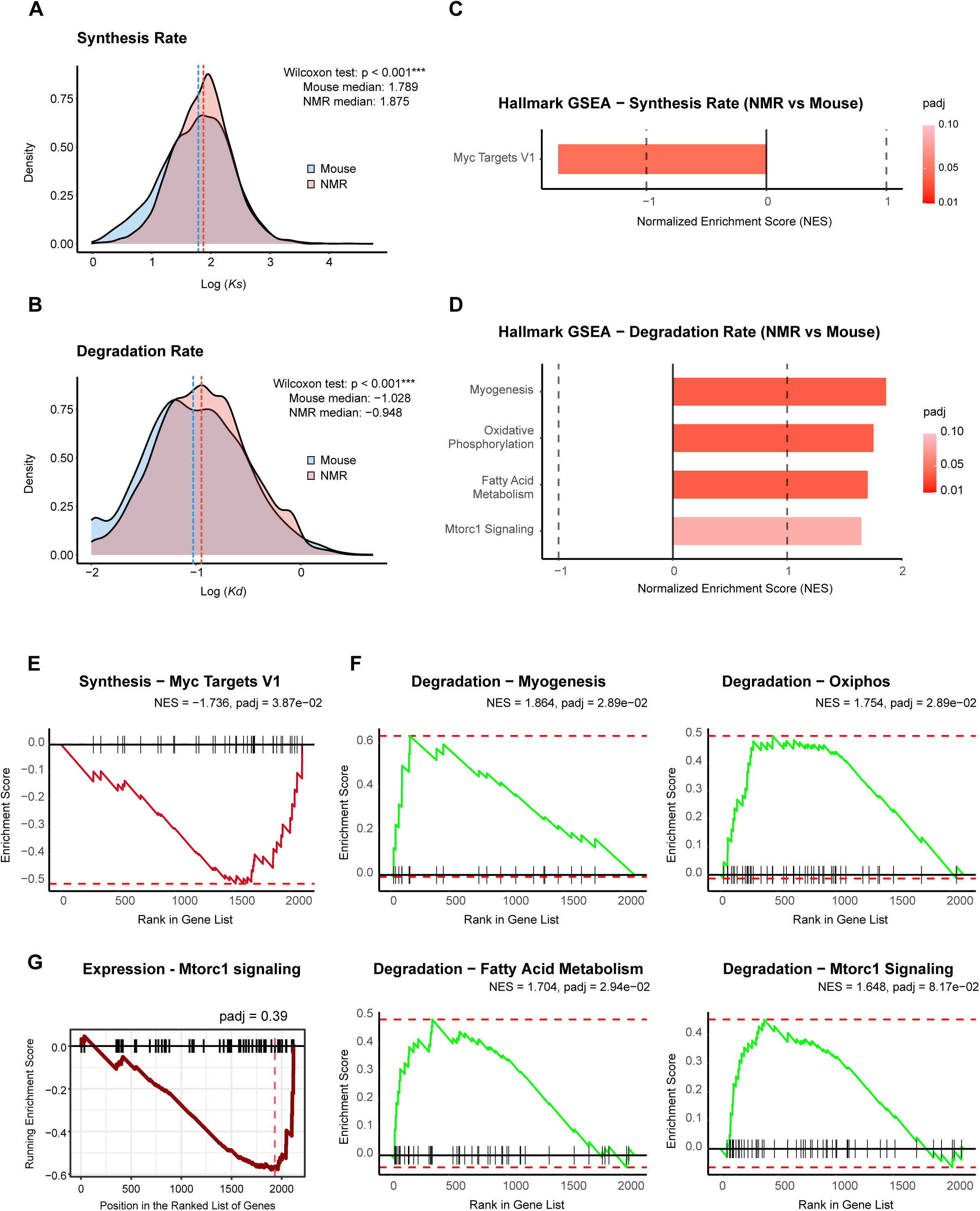
Comparison of NMR and mouse RNA kinetics. (A, B) RNA kinetics distribution of NMR and mouse genes. The common subset of genes (2,124 genes) was analyzed. RNA synthesis (A) and degradation (B) rates with the median in each animal (Red dashed line for NMR, blue for mouse). (C, D) Comparative RNA kinetics in NMR and mouse. The log_2_ fold change of each gene in *ks* (C) and *kd* (D) were subjected to GSEA with the mouse hallmark dataset. Adjusted *p*-value < 0.1 was used as the cutoff. (E, F) Enrichment plots of the GSEA in (C) and (D). (G) GSEA result of gene expression levels in the Hallmark MTORC1 pathway.

We next asked which pathways and/or cellular functions show altered RNA kinetics in NMR. To this end, we ordered genes by the fold changes of *ks* or *kd*, and ran gene set enrichment analysis (GSEA). Myc targets V1 was identified as the only dataset containing genes with altered *ks*, indicating that their synthesis rates are significantly lower in NMR than in mouse (Fig. 4C). GSEA of *kd* showed an enrichment of 4 pathways, all with faster degradation rates in NMR (Fig. 4D). The enrichment plots corresponding to these gene sets are shown in Fig. 4E-F.

One of these pathways is MTORC1, an established aging contributor which positively regulates aging (*i.e*., aging accelerator) (Mannick & Lamming, 2023; Ogawa et al., 2024; Panwar et al., 2023). Elevated *kd* of genes in MTORC1-pathway should result in reduced their gene expression, and these genes were indeed repressed in NMR, albeit the FDR was modest (Fig. 4G). Among genes, the biggest *kd* change was in *GRB10* (*kd* log_2_FC = 3.38). The protein GRB10 is a substrate of MTORC1, and is a key factor for crosstalk of nutrient sensing, aging and cognitive function (de Lucia et al., 2020). The difference in expression levels is similarly large (log_2_FC = −3.26) whereas that of the *ks* is marginal (−0.175), illustrating the influence of degradation on gene expression. Another finding was the elevated *kd* in the oxidative phosphorylation pathway (OXPHOS) and fatty acid metabolism of NMR. Consistent with the higher *kd*, a decrease in expression levels, though not significant, was also observed in these pathways in the GSEA (data not shown). Overall, the RNA kinetics of NMR show a distinct signature which influences gene expression levels.

### Aging-related genes may be subject to hidden controls by RNA turnover in NMR

As previously mentioned, gene expression is governed by the balance of *ks* and *kd* (*expression* = *ks/kd*) (Kawata et al., 2020). This means that the same expression level can be achieved by multiple combinations of *ks/kd* as long as the balance is maintained, even though the responsiveness of such genes can greatly differ (Fig. 5A). We next analyzed aging-related genes curated in the GenAge database (https://genomics.senescence.info/genes/index.html)(de Magalhães et al., 2005), and found several genes with similar expression levels but vastly different RNA kinetics (Fig. 5). Such genes include *APTX*, *NUDT1* (*MTH1*), and *XRCC5*, encoding DNA damage-repair proteins, the deficiency or overexpression of which has been linked to the alteration of aging and lifespan (Carroll et al., 2015; De Luca et al., 2013; Vogel et al., 1999). Despite little change in expression levels, the *ks* and *kd* are both elevated in these genes, suggesting that their RNA turnover and responsiveness are higher in NMR (Fig. 5B-D). Conversely, another DNA repair gene *FEN1* shows slower mRNA turnover, while the expression levels remain comparable (Fig. 5E). In summary, our data suggest that these genes are subject to distinct modes of expression control, despite the expression levels having barely changed.

**Figure 5.**
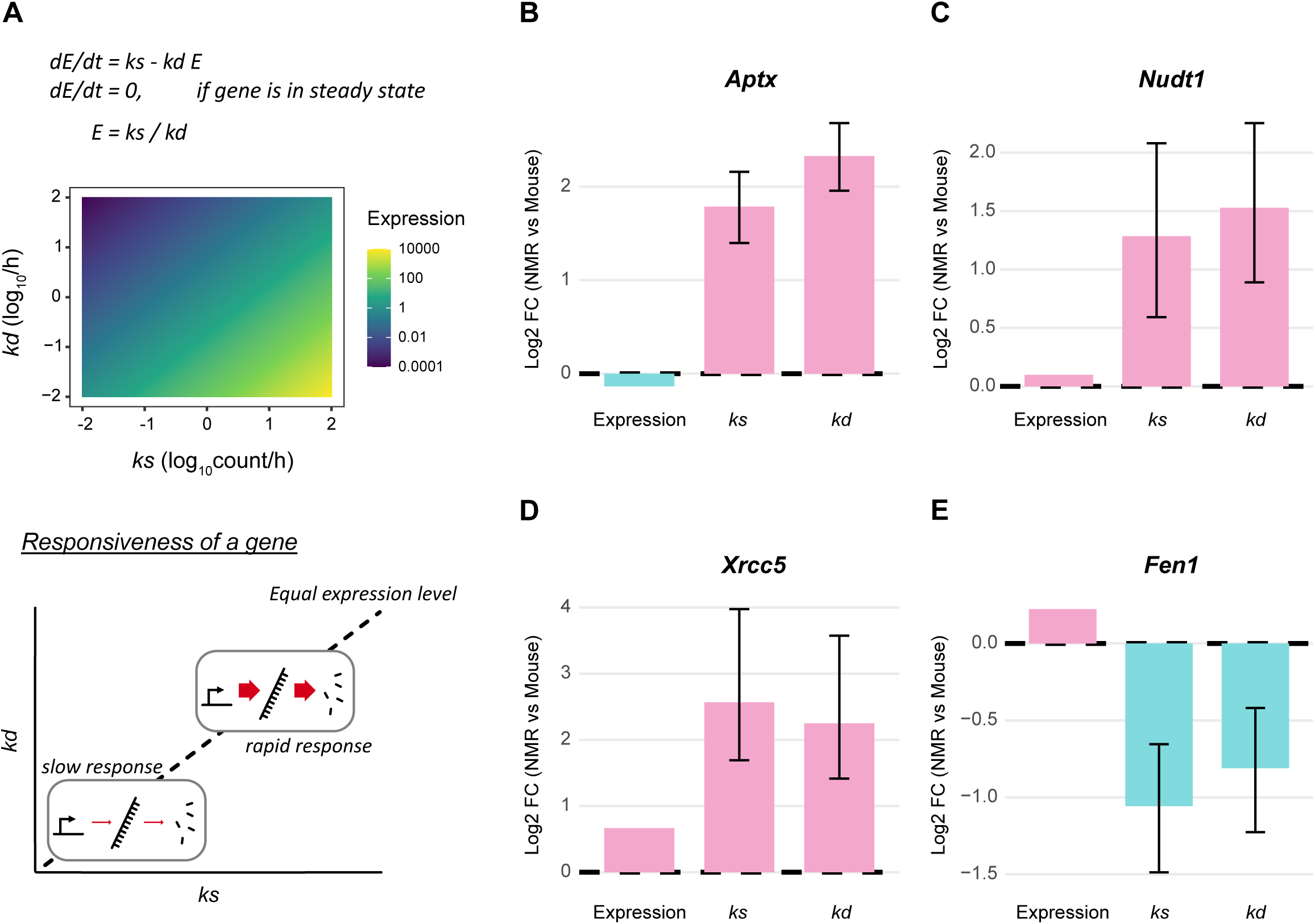
Aging-related genes with distinct RNA turnover but comparable gene expression levels. (A) Gene expression as a balance of RNA synthesis and degradation. Top: equations of gene expression and RNA kinetics. The equations were previously reported in (Kawata et al., 2020). E, expression level (e.g., read count). In steady state, the expression is defined by the ratio of *ks* and *kd*. Middle: simulated expression under various *ks* and *kd*. Note the unchanged expression along the diagonal (*ks/kd* unchanged). Bottom: biological interpretation of *ks, kd* and expression. With the same expression level, a gene may be transcribed into fast-turnover (rapid response) or stable (cost-saving) RNAs. (B-D) Examples of aging-related genes with elevated *ks* and *kd*, but with comparable expression levels. Each parameter was calculated as log_2_ fold change of NMR vs. mouse. (E) An example of aging-related genes with comparable expression level but reduced *ks and kd*. The error bars denote 95% confidence intervals, calculated by *grandR* (Rummel et al., 2023).

### Distinct control of RNA kinetics by transcript features in NMR

Our results have revealed the distinctive RNA kinetics of NMR fibroblasts and how these kinetics, especially the degradation as seen in the GSEA, are under unique control. To further elucidate the regulatory mechanisms, we investigated which transcript features influence RNA degradation by machine learning. Feature importance analysis via Random Forest and XGBoost revealed that the *kd* values of genes are in part explainable by the weighted feature-influences in both species (Random Forest, R^2^ ≈ 0.31 in NMR, 0.43 in mouse; XGBoost, R^2^ ≈ 0.30 in NMR, 0.45 in mouse) (Fig. 6A-B). Relative influence scores suggested overlapping yet distinct patterns of feature importance between NMR and mouse cells (Fig. 6C-D). In both species, the density of splice junctions (SJ) in the CDS was the most important feature. This is in alignment with a previous report predicting mRNA stability using a deep learning approach (Agarwal & Kelley, 2022). However, while the density of SJ in the overall mRNA was ranked second in NMR fibroblasts, the GC% of the CDS was ranked second in mouse cells. To further investigate this difference, we next attempted to directly explain the fold changes of *kd* (log_2_FC (*kd*)) between NMR and mouse fibroblasts. To this end, we used the difference in each feature (Δfeature) to explain the log_2_FC (*kd*) through machine learning. The results showed weaker model support (R^2^ ≈ 0.095 for Random Forest) (Fig. 6E); nevertheless, the CDS GC% was again the second-most important feature (Fig. 6F), a close match to the CDS A frequency, supporting the notion that nucleotide frequencies in the CDS govern RNA degradation to different degrees in NMR and mouse fibroblasts.

**Figure 6.**
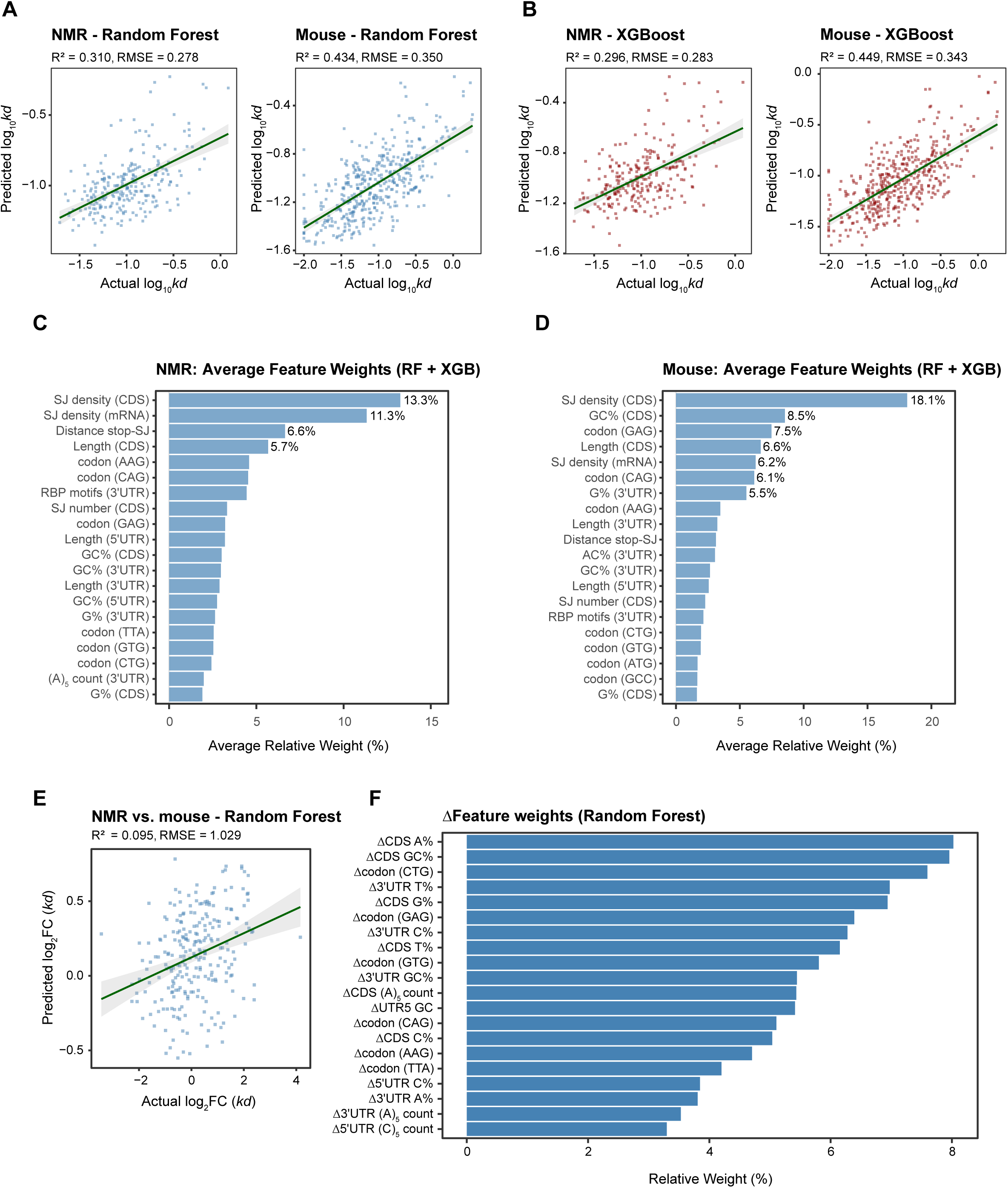
Feature weight analysis of RNA degradation rates. Transcript features and their influence on RNA degradation rates. (A, B) The actual and predicted log_10_*kd* in NMR and mouse by models trained by Random Forest (A) and XGBoost (B). RMSE, Root Mean Squared Error. (C) The relative feature weight for *kd* values in NMR. The top 20 features from the models in (A, B) are shown. The weight is the average of Random Forest and XGBoost models. Weights greater than 5% are indicated. RF, Random Forest; XGB, XGBoost. (D) The feature weight for *kd* values in mouse. (E) The actual and predicted log_2_FC of NMR and mouse *kd* by Random Forest. (F) The relative feature weight from the model in (D). The top 20 features are shown.

## Discussion

As a hub of cellular function, the transcriptome is recognized as a major player in aging research. Even though NMR has been extensively studied for its aging resistance, the underlying regulation of the transcriptome has remained an open question. In this study, we aimed to understand this regulatory mechanism in terms of the balance between RNA synthesis and degradation. We experimentally measured RNA kinetics in NMR for the first time and analyzed its significance in comparison to mice.

Certain genes in aging-associated pathway, including genes in the MTORC1 pathway, showed elevated degradation rates (larger *kd*) in NMR, even though their synthesis rates were not significantly altered. The most prominent example was *GRB10*, the substrate of MTORC1. The exact function of *GRB10* in aging remains elusive, but its deletion promotes self-renewal of mouse hematopoietic stem cells (Yan et al., 2016). It was also identified as an oncogene in glioma where the expression correlates with cell proliferation and cancer prognosis (Chen et al., 2022).

Therefore, the suppression of *GRB10* in NMR fibroblasts (log_2_FC = −3.26) may be beneficial for sustaining the cell population and maintaining the anti-oncogenic property of NMR. The down-regulation of *GRB10* was mostly explainable by the *kd* change (log_2_FC = 3.38, and the formula in Fig. 5A), highlighting the impact of RNA degradation on gene expression control in NMR. Other pathways with enhanced degradation were OXPHOS, and fatty acid metabolism which includes beta oxidation genes. In a previous study, the enzymatic activity of OXPHOS was found to be suppressed in NMR, and the oxygen consumption was lower than that in mouse at the mitochondrial, cellular, and organismal levels (Yap et al., 2022). Our finding of an elevated *kd* in mitochondrial respiration pathways, together with the reduced expression of those genes, aligns with the lower mitochondrial activity observed in NMR.

Our analysis of aging-related gene sets revealed that several genes, particularly genes involved in DNA damage repair, show an altered *ks/kd* balance even though their expression levels remain comparable. We previously showed, by theoretical and experimental analyses, that the swiftness of responses to environmental stimuli is determined predominantly by the RNA degradation rates (Kawata et al., 2020). Therefore, the alteration in *ks* and *kd* suggests that NMR fibroblasts may be more “ready” for DNA damage repair without changing the overall expression. Further supporting this notion, we previously detected a more rapid accumulation of 53BP1 and pATM foci (DNA damage response markers) in NMR neural stem/progenitor cells than in mouse cells (Yamamura et al., 2021). Even though this concept requires careful experimental validation, it is intriguing that the RNA turnover may act as a hidden layer of gene control in NMR and contribute to its stress response.

Another key question is which transcript feature enables the unique regulation of *kd* in NMR. The feature importance analysis revealed that the RNA kinetics of NMR and mouse cells are likely regulated by similar, though not identical features, including a variable influence of the CDS GC%. The CDS GC% is one of the strongest determinants of mRNA stability (Agarwal & Kelley, 2022) and the effect is in part explainable by RNA binding proteins (Jowhar et al., 2024), codon bias (Hia et al., 2019), and the subcellular localization of the RNA (Courel et al., 2019). Whether the observed *kd* difference in NMR is also influenced by such factors remains to be determined.

Among recent developments in RNA metabolic labeling, 4sU has become a *de facto* standard for its high labeling efficiency and low stress to the cell (Herzog et al., 2017; Kurosaki et al., 2021; Riml et al., 2017; Schofield et al., 2018). Nevertheless, 4sU inhibits rRNA synthesis, elicits a nucleolar stress response, and induces splicing defects (Altieri & Hertel, 2021; Burger et al., 2013). Through the development of the SLAM-seq analysis pipeline, Rummel et al. showed that 6.2% of the analyzed genes were strongly differentially expressed (Rummel et al., 2023). Therefore, a minor impact of 4sU on the transcriptome is inevitable, and computational methods to filter out such genes are important. We combined 3 intuitive criteria (Fig. 2A, B) to filter out not only DE genes but also genes with high variability. The 3-layer test is versatile and broadly applicable to RNA metabolic labeling experiments. Although we employed a simple ROC analysis and grid search, a future machine-learning approach may further improve the performance and robustness.

In conclusion, we successfully measured RNA synthesis and degradation kinetics in NMR and mouse cells. Altered RNA kinetics and the influence of distinct features in NMR cells hint at a unique transcriptome control. Our data will serve as a resource for the aging research community, as well as a starting point for future studies aimed at understanding aging-resistance mechanisms.

## Materials and Methods

### Animals

NMRs and mice were maintained in Kumamoto University as previously described (Kawamura et al., 2023). NMRs were raised in a stable condition with a temperature of 30°C ± 0.5°C and humidity of 55% ± 5%, in 12 h of light and dark cycles (Oka et al, 2022). C57BL/6N mice were purchased from CLEA Japan, kept at a temperature of 24.5°C ± 1.5°C and humidity of 50% ± 10%, with 12 h of light and dark cycles. All experiments were performed with approval of the Ethics Committees of Kumamoto University (approval No. A2020-042, A2022-079 and A2024-063) in compliance with *the Guide for the Care and Use of Laboratory Animals* (National Institutes of Health, Bethesda, MD).

### Cell culture

Primary adult skin fibroblasts were established from the back skin of a 1-year-old (young adult) male NMR, or a 6-week-old male C57BL/6N mouse, as previously described (Kawamura et al., 2023). Cells were maintained on gelatin-coated culture dishes (IWAKI) in Dulbecco’s modified Eagle’s medium (DMEM, Sigma) supplemented with 15% (for NMR) or 10% (mouse) fetal bovine serum (FBS; Nichirei), 1% Antibiotic-Antimycotic (Gibco), 2 mM L-glutamine (FUJIFILM Wako), and 0.1 mM nonessential amino acids (NEAA, FUJIFILM Wako). Passages and experiments were performed in an atmosphere-controlled chamber (SCI-tive DUAL hypoxia workstation, Baker Ruskinn) at 32°C with 5% CO_2_, 5% O_2_ and 90% N_2_. Fibroblasts were used for experiments within five passages.

### RNA metabolic labeling

The stock of 4-thiouridine (4sU, Toronto Research Chemicals) was prepared in ultrapure water (Invitrogen), and NaCl was added to adjust to an isotonic solution (285 mOsm/L ±10%). The medium containing 100 µM of 4sU was prepared a day before the experiment, and allowed to equilibrate in the culture chamber overnight. Cells in mid log-phase were labelled with 4sU by exchanging the medium with 4sU-containing medium and incubated for up to 8 hours to allow 4sU incorporation into new RNA before harvest (Fig. 1B). At each time point, the cells were washed with prewarmed PBS (-), lysed in 3 mL of RNAiso Plus (Takara Bio) and kept at −80 ⁰C until use.

### RNA extraction and nucleotide conversion

RNA in RNAiso Plus was extracted by following the manufacturer’s instruction. After its quality and quantity were confirmed by Nanodrop One (Thermo Fisher Scientific) and Bioanalyzer 2100 with the RNA nano 6000 kit (Agilent Technologies), the incorporated 4sU was converted to cytosine following the TUC-seq DUAL protocol (Gasser et al., 2020) with minor modifications (Fig. 1C). Briefly, 4 µg of total RNA was incubated with 0.45 mM osmium tetroxide (OsO_4_; FUJIFILM Wako) and 180 mM of NH_4_Cl-NH_4_OH buffer (pH 8.88) at 40 ⁰C for 2 hours with agitation at 1,200 rpm. Osmium was removed by washing three times with ultrapure water on an Amicon Ultra centrifugal concentrator (MWCO 3k; Merck Millipore), and the RNA was further incubated in 125 mM Hydrazine-H_2_O in TE buffer (pH: 8.11; 125 mM Tris, 1.25 mM EDTA) at 40 ⁰C for 2 hours with agitation at 1,200 rpm. The RNA was subsequently recovered using the RNA Clean-Up kit (Takara Bio). Before the library preparation, the RNA was treated with DNase I (Takara Bio) to ensure the removal of residual genomic DNA. For sequencing, libraries were prepared with the DNBSEQ Eukaryotic Strand-specific mRNA library kit (PE150) and analyzed on the DNBSEQ sequencing platform. Library preparation and sequencing were performed by BGI Japan, and a minimum of 20 million read pairs per RNA sample were obtained.

### Calculation of RNA kinetics

Reads were filtered by fastp ver. 0.23.2 with default parameters, aligned to the reference genome using STAR ver. 2.7.10a (Dobin et al., 2013) with options ‘--outFilterMultimapNmax 1 --outSAMattributes MD NH nM --chimSegmentMin 10 --alignTranscriptsPerReadNmax 100000 --outFilterMismatchNmax 20’. The following genome and annotation files were obtained from Ensembl release 114: NMR, Heterocephalus_glaber_female.Naked_mole-rat_maternal.dna_sm.primary_assembly.fa.gz (1-29, MT, X) and the corresponding annotation Heterocephalus_glaber_female.Naked_mole-rat_maternal.114.chr.gtf.gz; Mouse, Mus_musculus.GRCm39.114.chr.gtf.gz and the corresponding annotation Mus_musculus.GRCm39.dna_sm.primary_assembly.fa.gz. After mapping, the proportion of 4sU-labeled RNA at each time point was estimated by Bayesian network analysis using GRAND-SLAM ver. 2.0.7b (Jürges et al., 2018) and the RNA kinetics calculated by the accompanying R package *grandR* ver. 0.2.6 (Rummel et al., 2023). After the initial kinetics-fitting by *grandR*, the kinetics estimation was further polished by taking the different 4sU incorporation rates between species into account using the *grandR* function *CalibrateEffectiveLabelingTimeKineticFit* (Rummel et al., 2023). For differential gene expression analysis, the mapped read counts were extracted from the GRAND-SLAM output and analyzed by *DESeq2* size factor normalization (Love et al., 2014). For the calculation of *ks*, the DESeq2-normalized read count was used as the expression level.

### Detection of steady-state genes by 3-layer test

Steady-state genes were identified using a three-layer approach, in which three complementary criteria were applied: 1) the maximum fold changes between any of the two time points are smaller than the threshold; 2) coefficient of deviation within a time point (or (max - mean)/mean for 2 replicates) is smaller than the threshold for any of the time points; 3) the null hypothesis is not rejected in a differential expression analysis (*e.g*., DESeq2). For the 3rd criteria, we chose a likelihood ratio test (LRT) using DESeq2 (ver. 1.46.0) (Love et al., 2014) with a reduced (∼1) versus a full model (∼timepoint), and genes with significance were assigned non-stable. To optimize threshold parameters, we prepared a gold standard dataset by randomly selecting 500 genes from our other transcriptome experiment (Matsubara et al., unpublished data) and manually classifying their expression patterns as steady (TRUE) or unsteady (FALSE), based on visual assessment of expression profiles (Fig. 2A-B and supplemental Fig. 1). Optimal thresholds were then determined using *ROCR* v1.0-11 (Sing et al., 2005) to depict the ROC curve followed by a grid search with Youden index maximization (*Caret*, ver. 7.0-1: https://cran.r-project.org/package=caret). Grid search evaluated parameters ranging from 1.00 to 1.50 for fold-change assessment; from 0.10 to 0.50 for variability thresholds; from 0.01 to 0.50 for DESeq2 p.adj (each with 0.01 increments).

After establishing the parameters, all genes were classified according to their expression profiles. Genes were classified as "steady" only when they satisfied all three criteria, whereas those failing any of the test layers were classified as "unsteady".

### Gene ontology term enrichment analysis (GO analysis)

GO analysis was performed using the g:Profiler online server (https://biit.cs.ut.ee/gprofiler/gost) (Kolberg et al., 2023). The *ks* and *kd* values in NMR RNA (7,281 genes) were binned by the 1^st^ and 3^rd^ quantiles, and genes categorized in 9 subgroups from A to I (Fig. 3A). Genes in each category (excluding the group E) were analyzed for GO-term enrichment using all 7,281 genes as background.

### Gene set enrichment analysis (GSEA)

GSEA (Mootha et al., 2003; Subramanian et al., 2005) was performed using the R *fgsea* package ver. 1.32.4 (Sergushichev, 2016) to identify significantly enriched biological pathways and functional categories. Gene sets were obtained from the Molecular Signatures Database (MSigDB) using the R *msigdbr* package ver. 24.1.0, including Hallmark gene sets and Gene Ontology biological processes (Castanza et al., 2023). Pre-ranked gene lists were generated based on log2 fold changes from differential expression analysis. Pathways with p-adjusted < 0.05 were considered significantly enriched.

### Feature importance analysis

Features were extracted from the transcripts with the “Ensembl_canonical” tag. For NMR, UTR sequences were extracted from RefSeq genome and annotation (GCF_000247695.1_HetGla_female_1.0_genomic.fna/gtf) and integrated with the feature table made from Ensembl. Then, the *kd* values and the extracted transcript features were used to train Random Forest and XGBoost models by using the R packages *randomForest* and *xgboost* (https://CRAN.R-project.org/package=randomForest; https://cran.r-project.org/package=xgboost). The model performance was assessed by R^2^ of actual and predicted *kd* values from the test data subset. To identify features defining the difference in RNA kinetics, kd fold changes (log_2_FC *kd*) and features differences (ΔFeatures) were used for training.

## Data availability

The RNA-seq data for NMR and mouse 4sU-labeling experiments are available in DDBJ Sequence Read Archive with the accession numbers DRR797616-DRR797639, under BioProject (PRJDB39662). BioSample metadata are available with the accession numbers SAMD01779538-SAMD01779561.

## Acknowledgments

We thank Dr. Nobuhito Goda (Waseda University, Tokyo, Japan) for kindly letting us use the hypoxia workstation. We also thank Dr. Therese Solberg (Chiba University, Chiba, Japan, and Keio University, Tokyo, Japan) and Toson Hamza (the University of Tokyo, Japan) for valuable comments and grammatical advises. Computational analyses were performed in part on the NIG supercomputer at ROIS National Institute of Genetics, Japan. J.W. also acknowledges *the University of Tokyo Foundation for Supporting International Students Scholarship* for personal financial support during the master’s program.

## Author contribution statement

R.M., Conceptualization, Investigation, Formal analysis, Visualization, Funding acquisition, Writing – original draft, Writing – review & editing. W.J., Formal analysis, Visualization. A.N.O., Investigation. Y.K., Investigation, Funding acquisition, Writing – review & editing. M.H., Supervision, Writing – review & editing. K.M., Investigation, Funding acquisition, Writing – review & editing. N.A., Conceptualization, Supervision, Funding acquisition, Writing – review & editing.

## Supplemental figures and tables

**Supplemental Figure 1.** The manually-curated ground truth of gene expression profiles used for the 3-layer test, related to Fig. 2.

**Supplemental Figure 2.** Detailed schematic representation of the 3-layer test, related to Fig. 2. (top) an example of RNA expression counts over a time series. LOESS, locally estimated scatterplot smoothing. (i-iii) the three criteria composing 3-layer test. The thresholds (threshold_i-iii_) were predetermined from a ground truth data set (see methods). (i) Significance in the differential expression analysis (*e.g.*, DESeq2) with a reduced (∼1) versus a full model (∼timepoint). The small box on the right denotes the reduced model in which a temporal parameter is not considered. (ii) The maximum value of replicate differences within each time point (*d*_1_ will be selected in this example). (iii) The maximum of the fold-changes between any combinations of the time points. The dashed lines denote the highest and lowest expression values in the time course.

**Supplemental Table 1.** The RNA kinetics of NMR and mouse fibroblasts.

**Supplemental Table 2.** GO-term enrichment of NMR genes in different RNA kinetics. Each Excel tab represents the enriched terms in the category defined in Fig. 3. The category A yielded no significant enrichment.

