## Supplemental Figure 1 for "RNA degradation modulates unique aging-related gene expression in naked mole-rats"

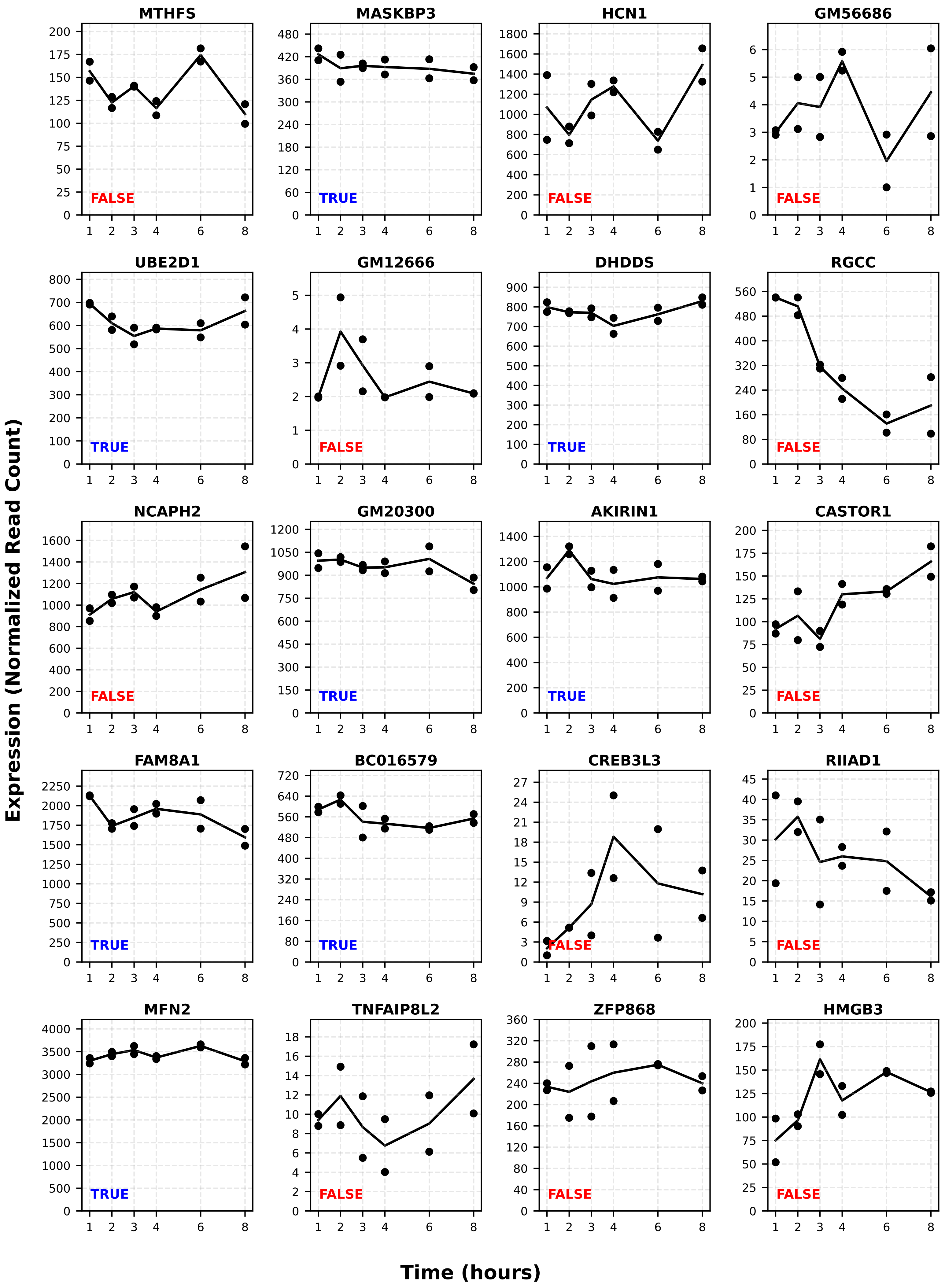
MTHFS
MASKBP3
HCN1
GM56686
UBE2D1
GM12666
DHDDS
RGCC
NCAPH2
GM20300
AKIRIN1
CASTOR1
FAM8A1
BC016579
CREB3L3
RIIAD1
MFN2
TNFAIP8L2
ZFP868
HMGB3
FALSE
TRUE
Expression (Normalized Read Count)
Time (hours)

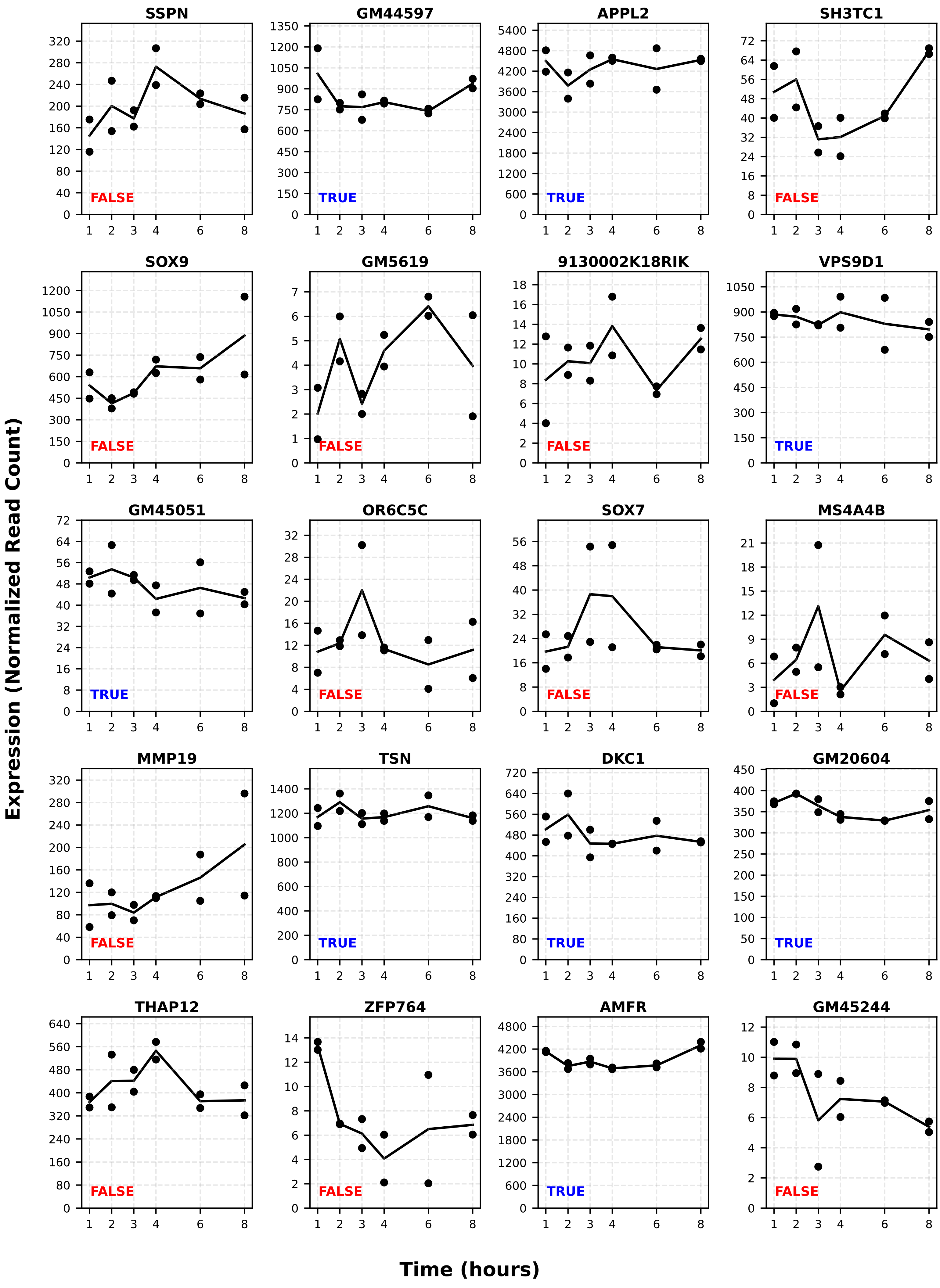
SSPN
FALSE
GM44597
TRUE
APPL2
TRUE
SH3TC1
FALSE
SOX9
FALSE
GM5619
FALSE
9130002K18RIK
FALSE
VPS9D1
TRUE
GM45051
TRUE
OR6C5C
FALSE
SOX7
FALSE
MS4A4B
FALSE
MMP19
FALSE
TSN
TRUE
DKC1
TRUE
GM20604
TRUE
THAP12
FALSE
ZFP764
FALSE
AMFR
TRUE
GM45244
FALSE
Expression (Normalized Read Count)
Time (hours)

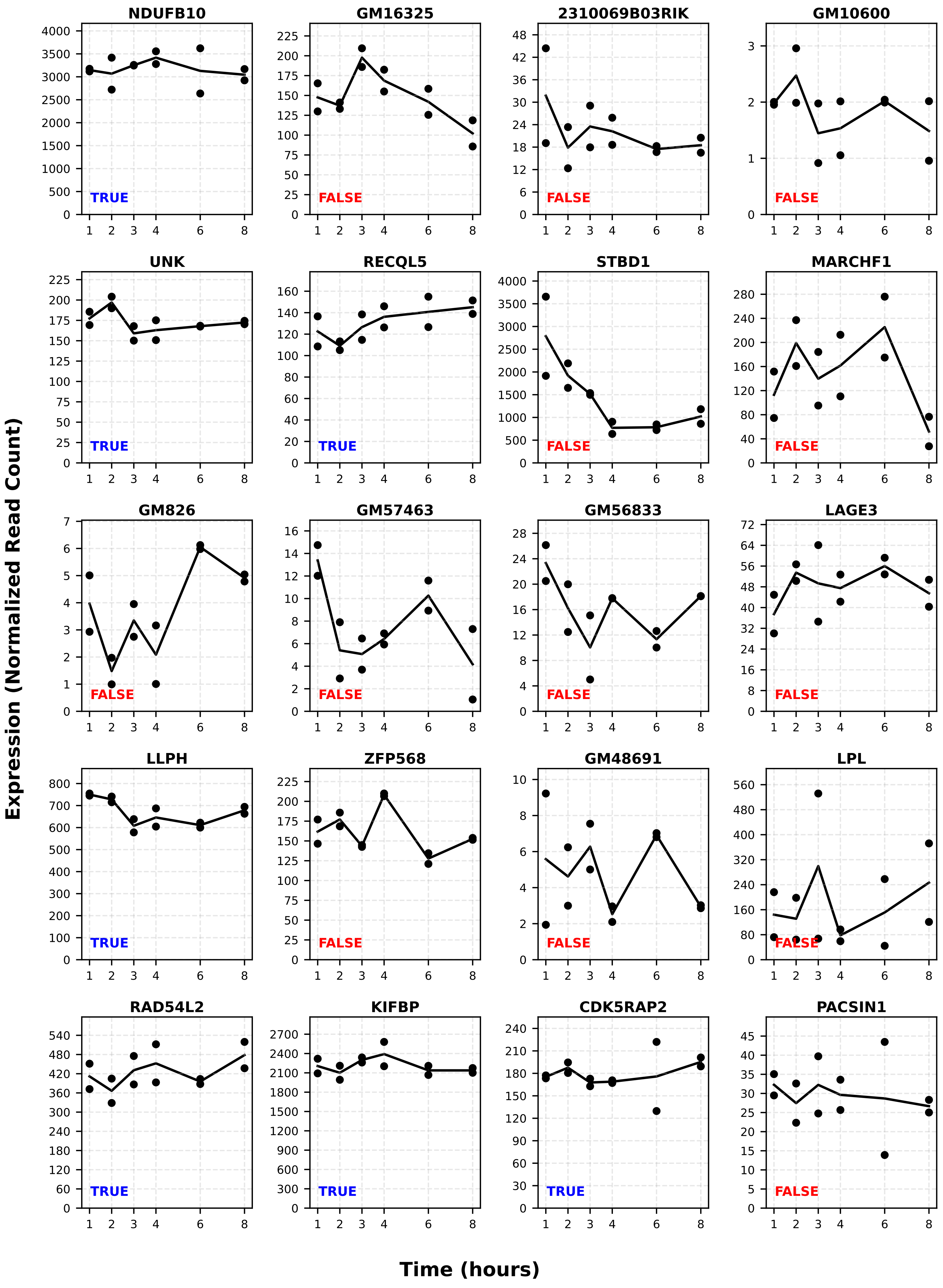
NDUFB10
TRUE
GM16325
FALSE
2310069B03RIK
FALSE
GM10600
FALSE
UNK
TRUE
RECQL5
TRUE
STBD1
FALSE
MARCHF1
FALSE
GM826
FALSE
GM57463
FALSE
GM56833
FALSE
LAGE3
FALSE
LLPH
TRUE
ZFP568
FALSE
GM48691
FALSE
LPL
FALSE
RAD54L2
TRUE
KIFBP
TRUE
CDK5RAP2
TRUE
PACSIN1
FALSE
Expression (Normalized Read Count)
Time (hours)

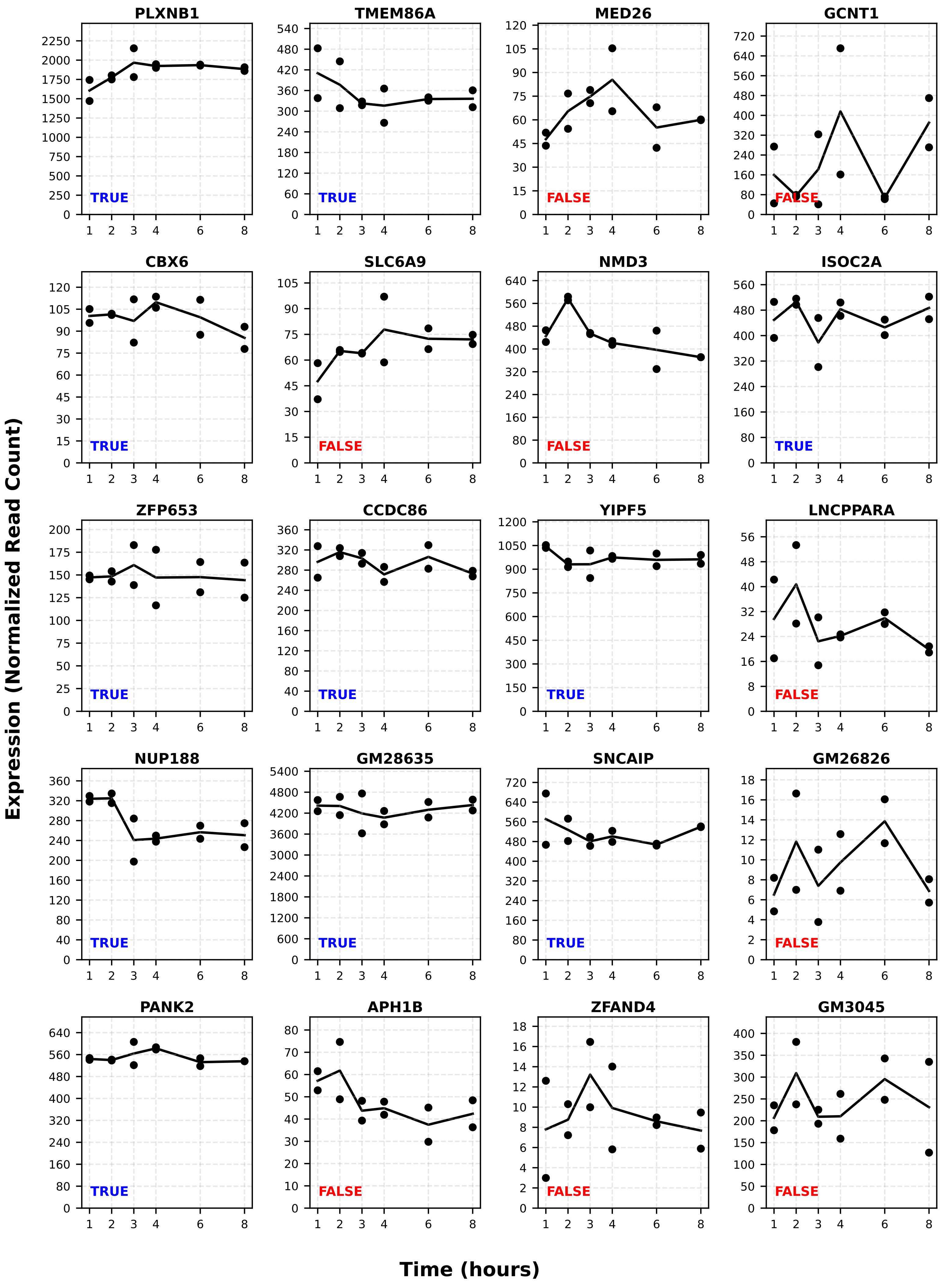

PLXNB1
TMEM86A
MED26
GCNT1
CBX6
SLC6A9
NMD3
ISOC2A
ZFP653
CCDC86
YIPF5
LNCPPARA
NUP188
GM28635
SNCAIP
GM26826
PANK2
APH1B
ZFAND4
GM3045
TRUE
FALSE
Expression (Normalized Read Count)
Time (hours)

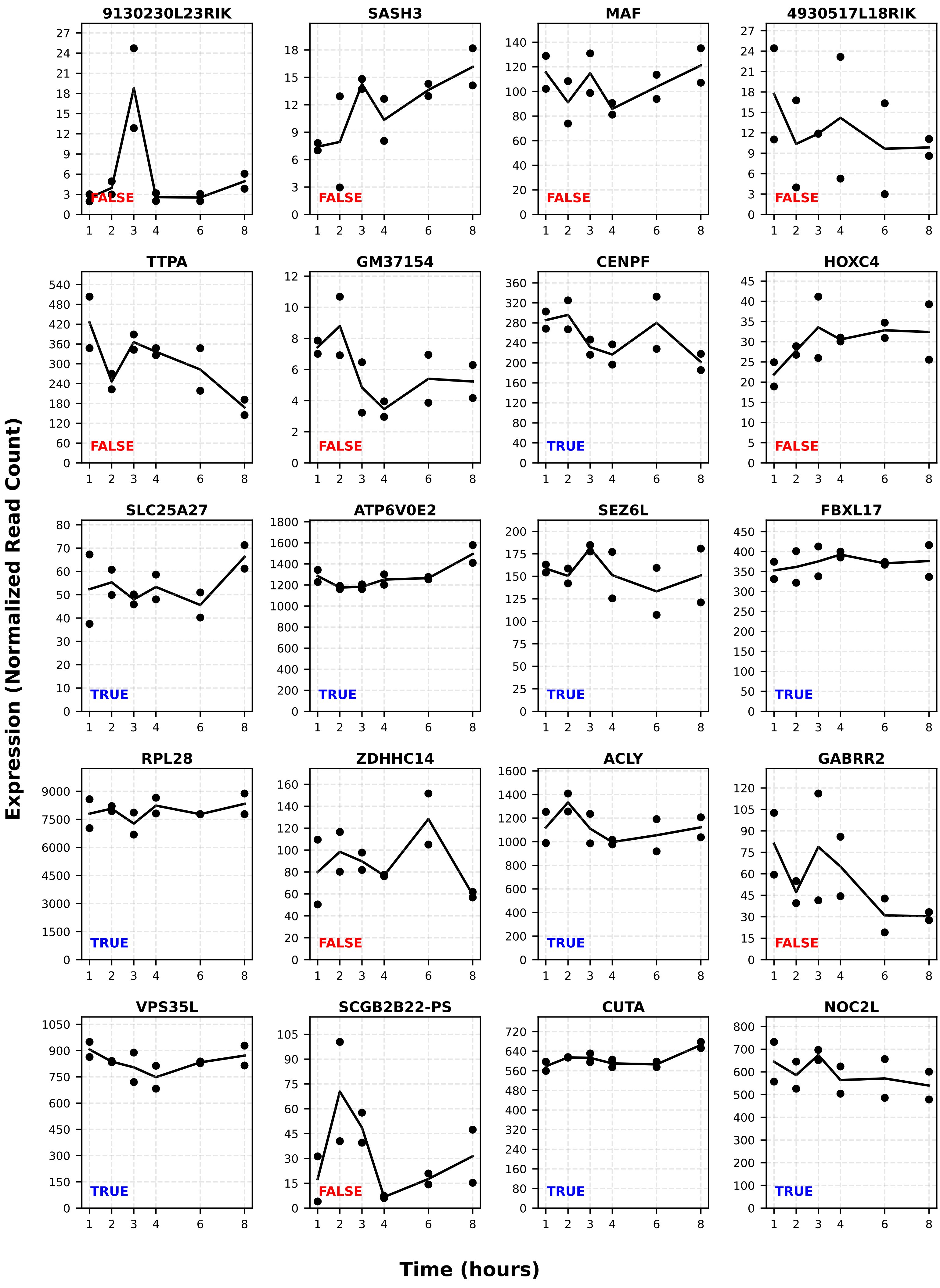

9130230L23RIK
SASH3
MAF
4930517L18RIK
TTPA
GM37154
CENPF
HOXC4
SLC25A27
ATP6V0E2
SEZ6L
FBXL17
RPL28
ZDHHC14
ACLY
GABRR2
VPS35L
SCGB2B22-PS
CUTA
NOC2L
FALSE
TRUE
Expression (Normalized Read Count)
Time (hours)

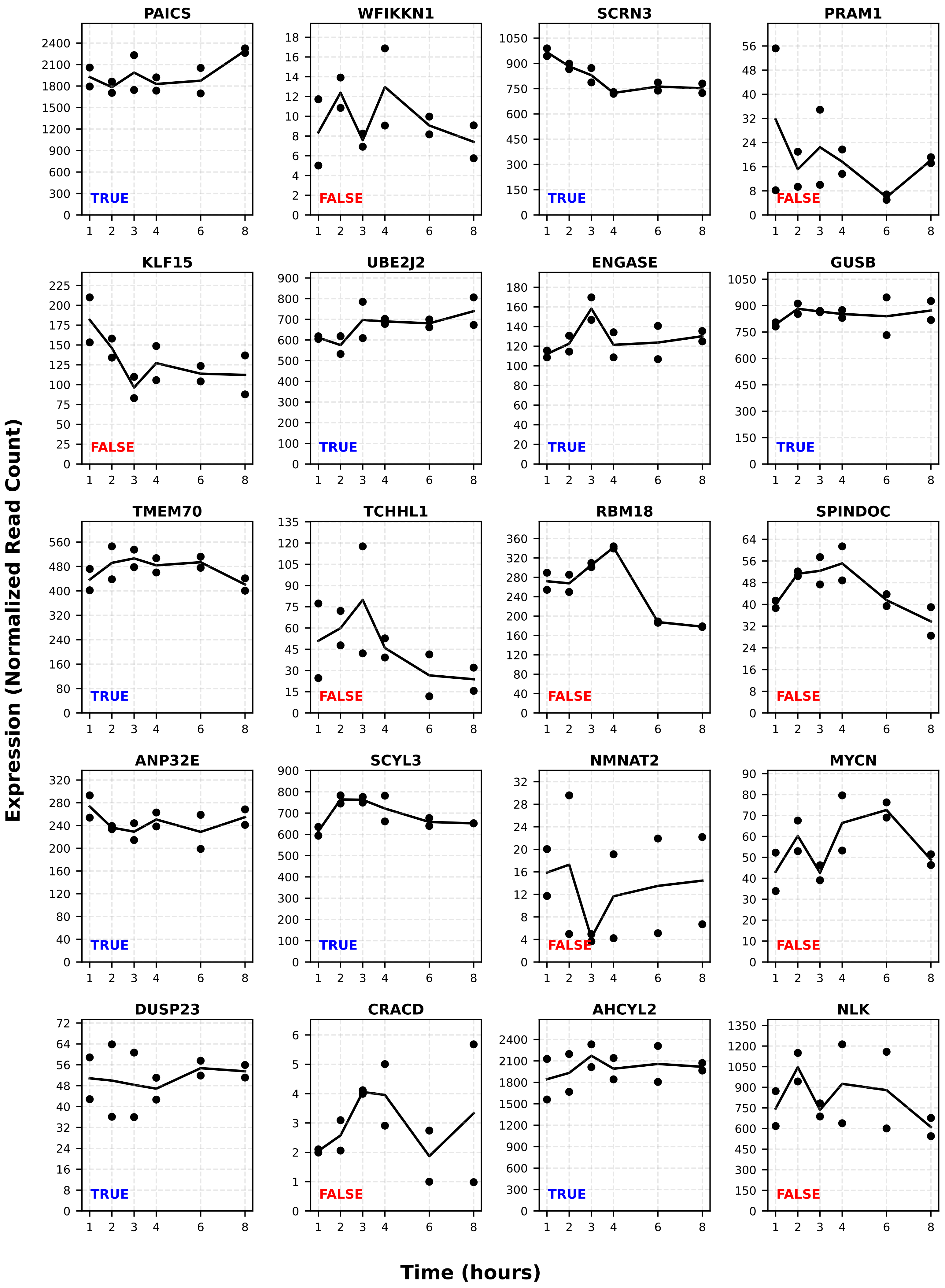

PAICS
WFIKKN1
SCRN3
PRAM1
KLF15
UBE2J2
ENGASE
GUSB
TMEM70
TCHHL1
RBM18
SPINDOC
ANP32E
SCYL3
NMNAT2
MYCN
DUSP23
CRACD
AHCYL2
NLK
TRUE
FALSE
Expression (Normalized Read Count)
Time (hours)

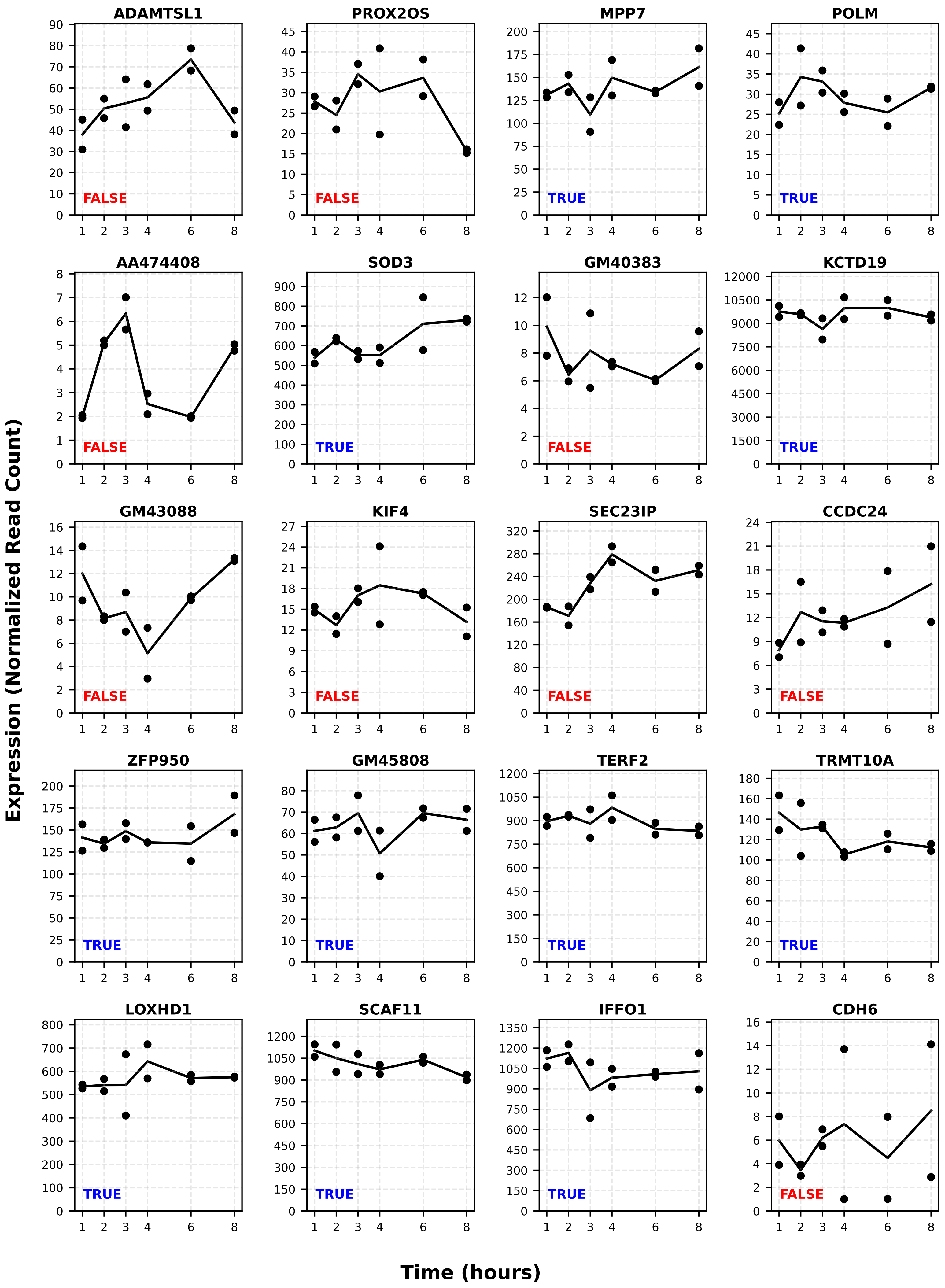
ADAMTSL1
FALSE
PROX2OS
FALSE
MPP7
TRUE
POLM
TRUE
AA474408
FALSE
SOD3
TRUE
GM40383
FALSE
KCTD19
TRUE
GM43088
FALSE
KIF4
FALSE
SEC23IP
FALSE
CCDC24
FALSE
ZFP950
TRUE
GM45808
TRUE
TERF2
TRUE
TRMT10A
TRUE
LOXHD1
TRUE
SCAF11
TRUE
IFFO1
TRUE
CDH6
FALSE
Expression (Normalized Read Count)
Time (hours)

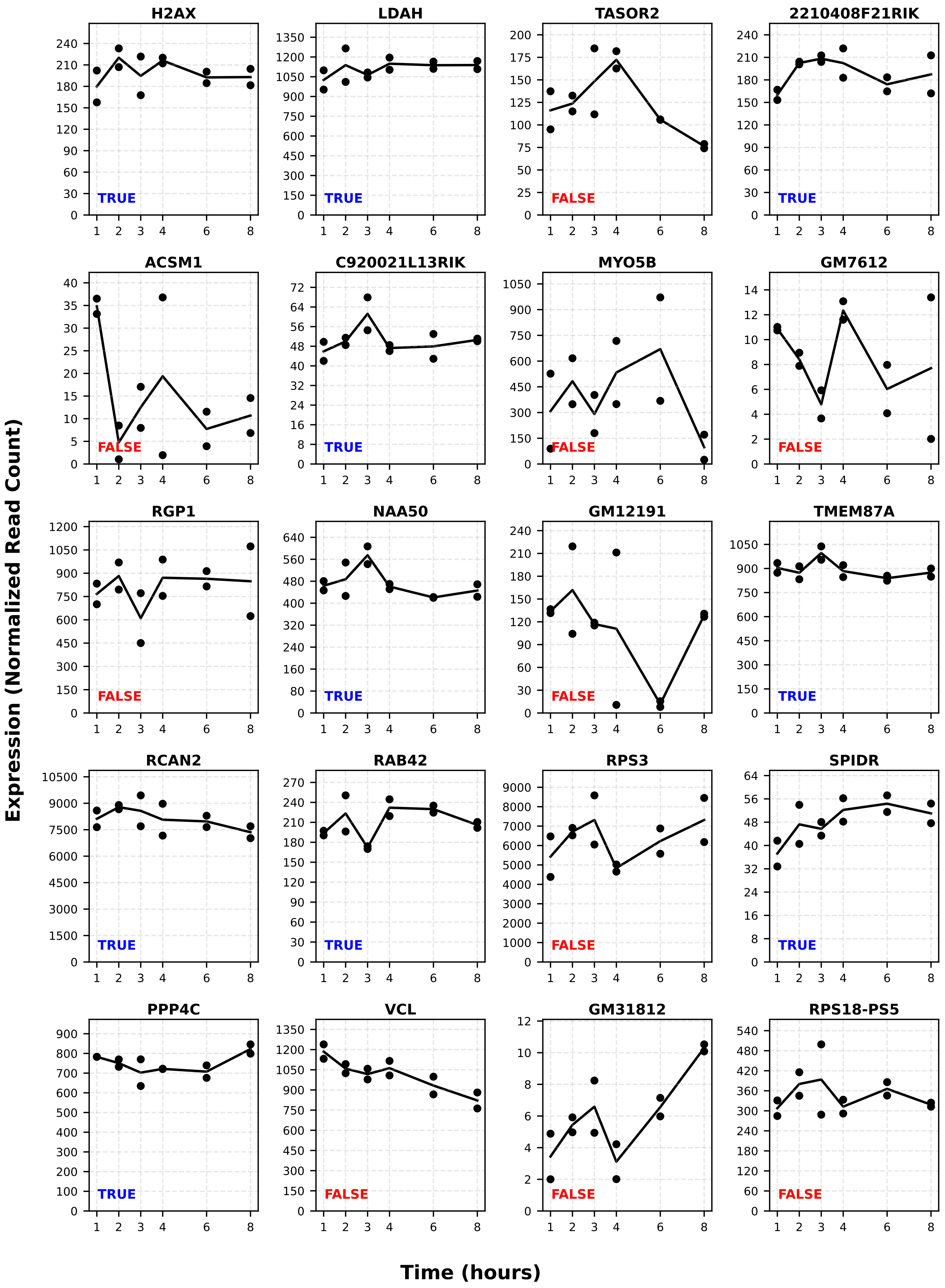
H2AX
TRUE
LDAH
TRUE
TASOR2
FALSE
2210408F21RIK
TRUE
ACSM1
FALSE
C920021L13RIK
TRUE
MYO5B
FALSE
GM7612
FALSE
RGP1
FALSE
NAA50
TRUE
GM12191
FALSE
TMEM87A
TRUE
RCAN2
TRUE
RAB42
TRUE
RPS3
FALSE
SPIDR
TRUE
PPP4C
TRUE
VCL
FALSE
GM31812
FALSE
RPS18-PS5
FALSE
Expression (Normalized Read Count)
Time (hours)

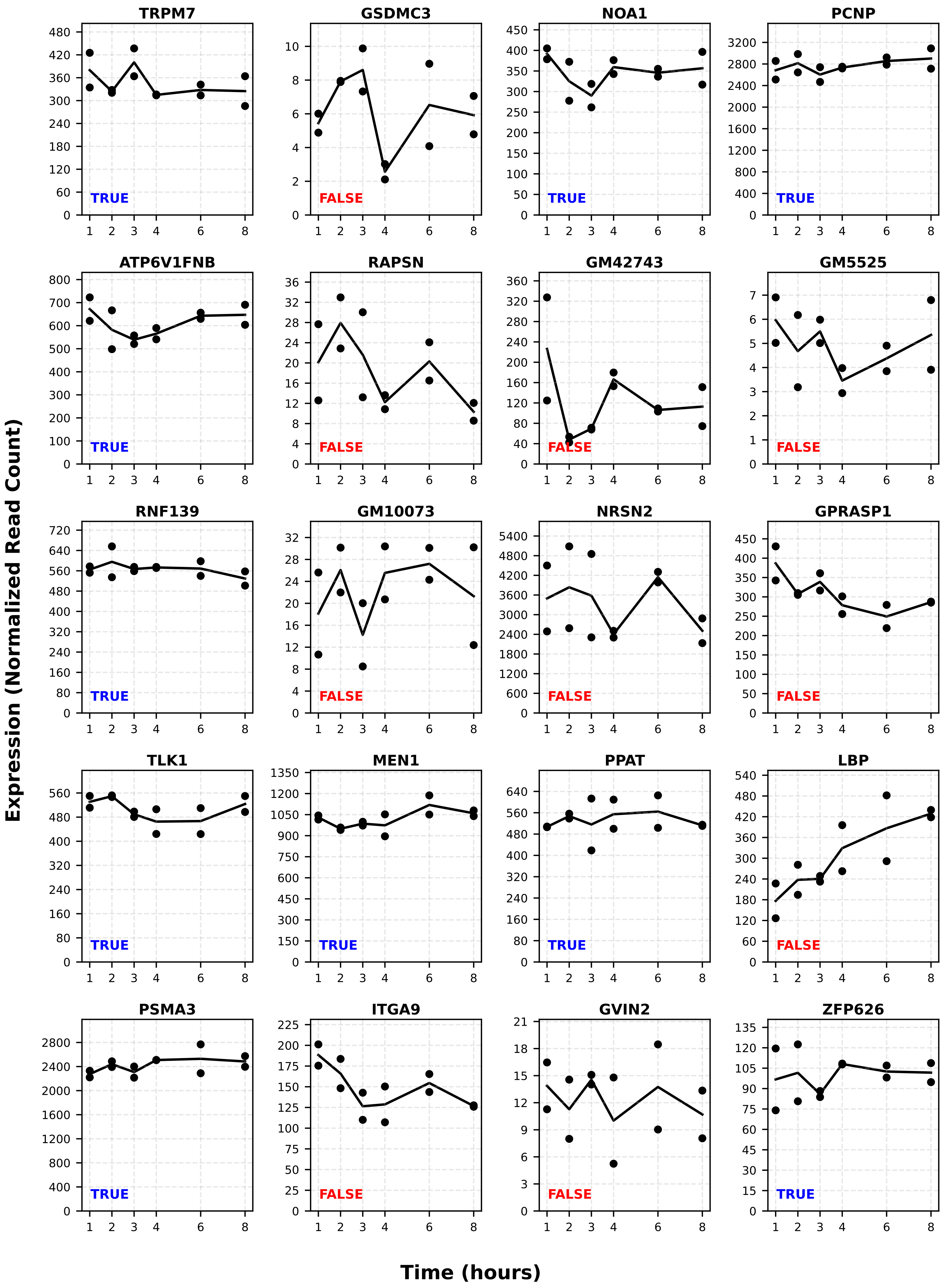

TRPM7
TRUE
GSDMC3
FALSE
NOA1
TRUE
PCNP
TRUE
ATP6V1FNB
TRUE
RAPSN
FALSE
GM42743
FALSE
GM5525
FALSE
RNF139
TRUE
GM10073
FALSE
NRSN2
FALSE
GPRASP1
FALSE
TLK1
TRUE
MEN1
TRUE
PPAT
TRUE
LBP
FALSE
PSMA3
TRUE
ITGA9
FALSE
GVIN2
FALSE
ZFP626
TRUE
Expression (Normalized Read Count)
Time (hours)

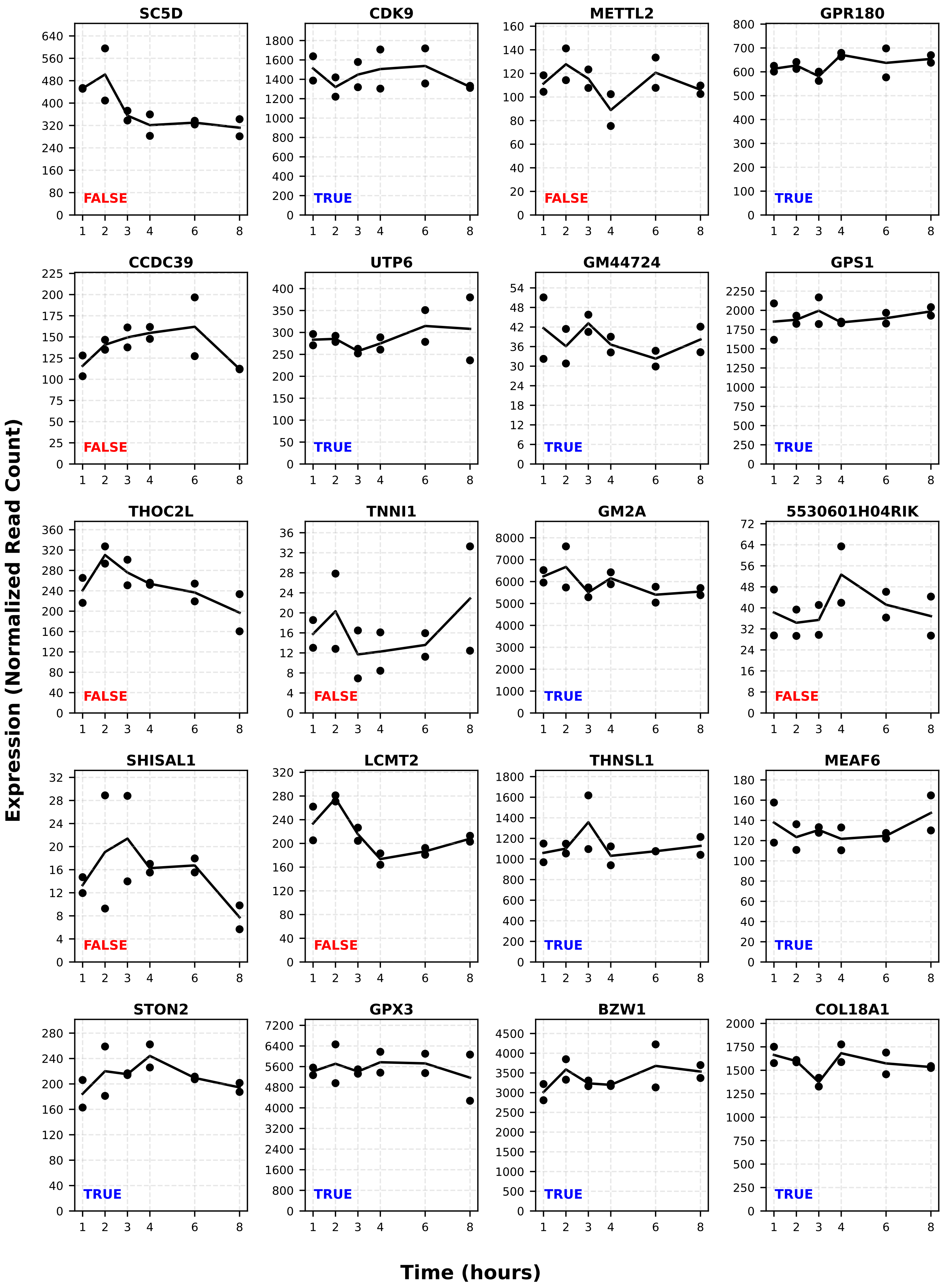
SC5D
FALSE
CDK9
TRUE
METTL2
FALSE
GPR180
TRUE
CCDC39
FALSE
UTP6
TRUE
GM44724
TRUE
GPS1
TRUE
THOC2L
FALSE
TNNI1
FALSE
GM2A
TRUE
5530601H04RIK
FALSE
SHISAL1
FALSE
LCMT2
FALSE
THNSL1
TRUE
MEAF6
TRUE
STON2
TRUE
GPX3
TRUE
BZW1
TRUE
COL18A1
TRUE
Expression (Normalized Read Count)
Time (hours)

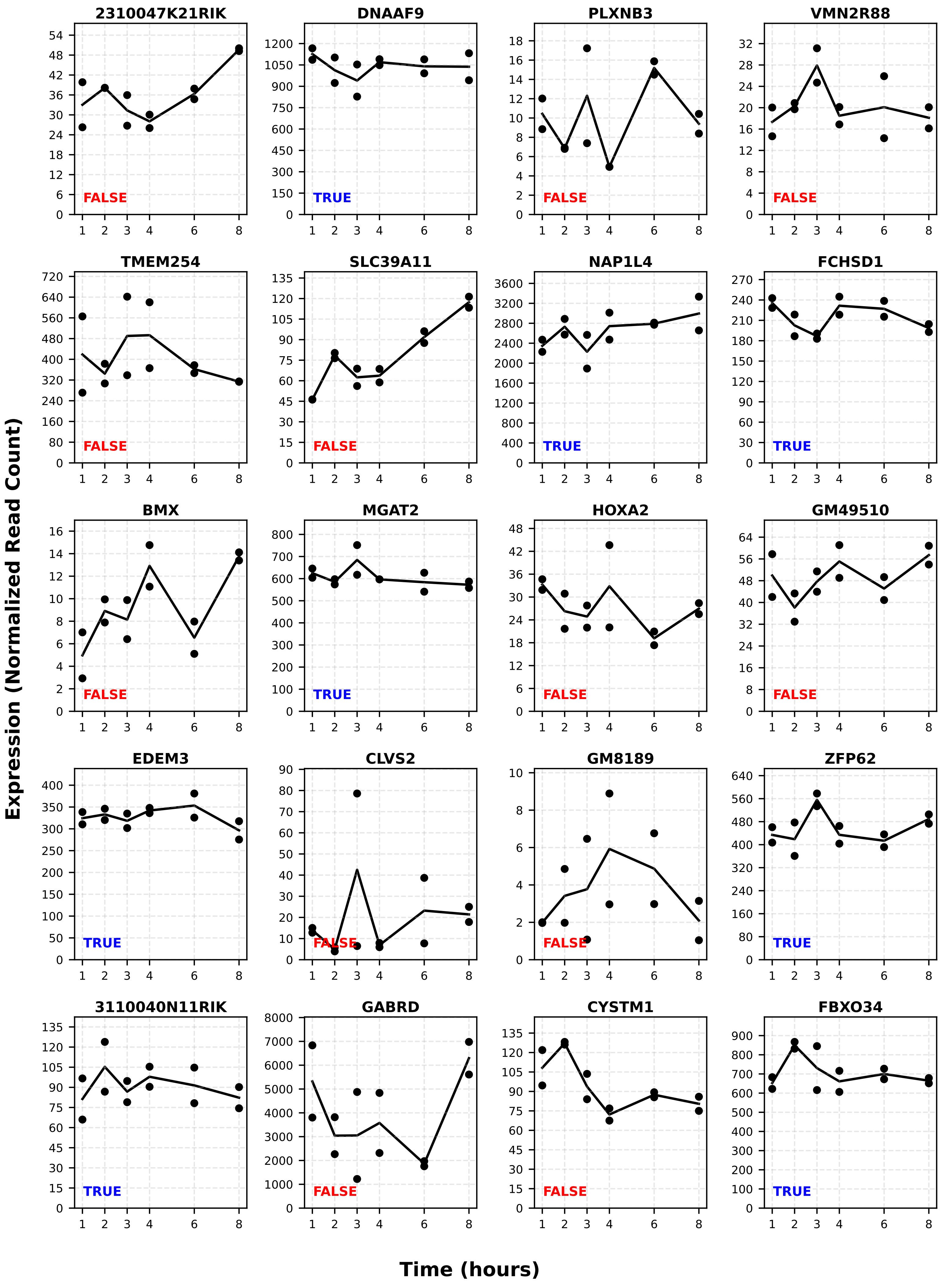
2310047K21RIK
FALSE
DNAAF9
TRUE
PLXNB3
FALSE
VMN2R88
FALSE
TMEM254
FALSE
SLC39A11
FALSE
NAP1L4
TRUE
FCHSD1
TRUE
BMX
FALSE
MGAT2
TRUE
HOXA2
FALSE
GM49510
FALSE
EDEM3
TRUE
CLVS2
FALSE
GM8189
FALSE
ZFP62
TRUE
3110040N11RIK
TRUE
GABRD
FALSE
CYSTM1
FALSE
FBXO34
TRUE
Expression (Normalized Read Count)
Time (hours)

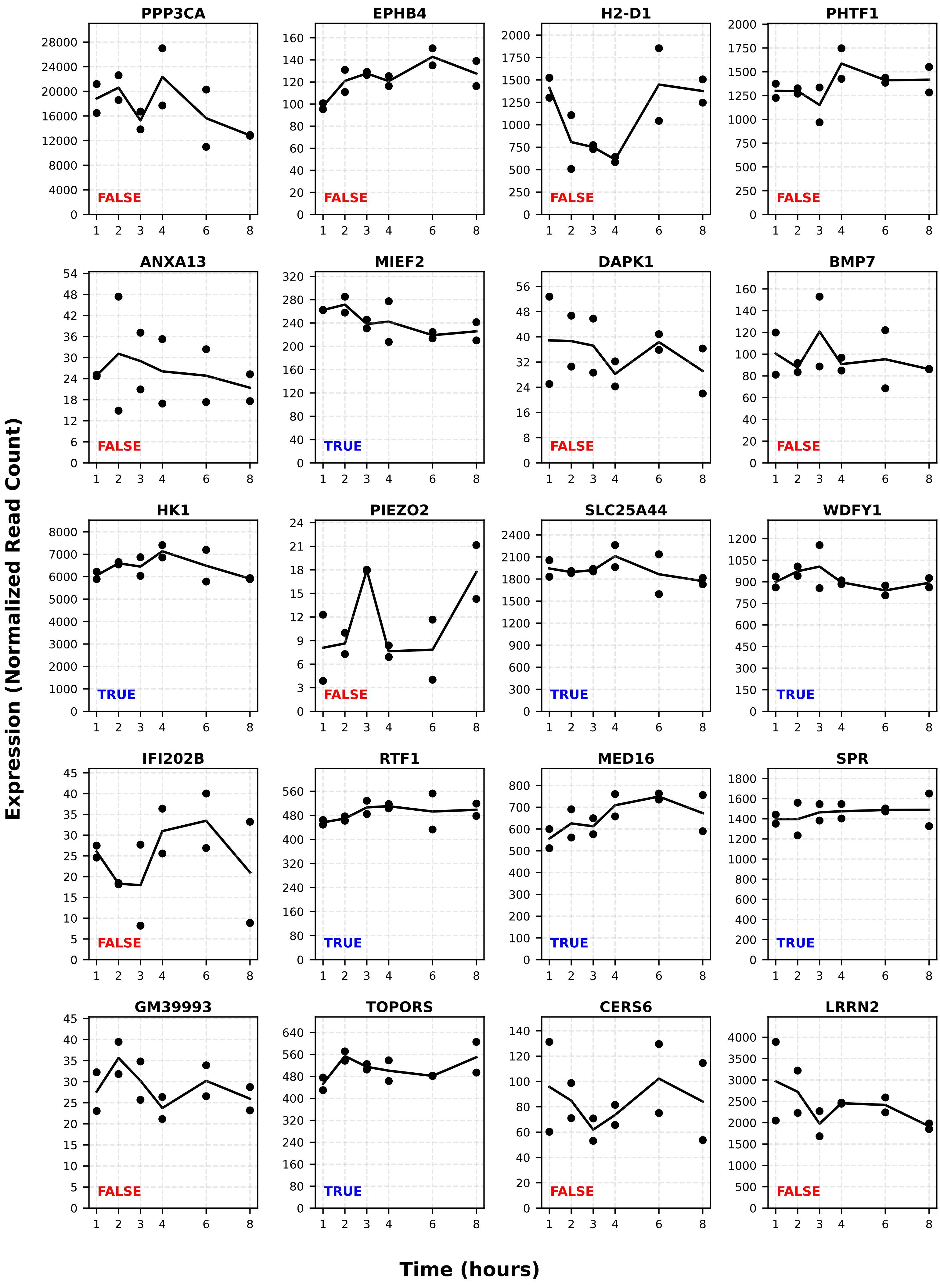

PPP3CA
FALSE
EPHB4
FALSE
H2-D1
FALSE
PHTF1
FALSE
ANXA13
FALSE
MIEF2
TRUE
DAPK1
FALSE
BMP7
FALSE
HK1
TRUE
PIEZO2
FALSE
SLC25A44
TRUE
WDFY1
TRUE
IFI202B
FALSE
RTF1
TRUE
MED16
TRUE
SPR
TRUE
GM39993
FALSE
TOPORS
TRUE
CERS6
FALSE
LRRN2
FALSE
Expression (Normalized Read Count)
Time (hours)

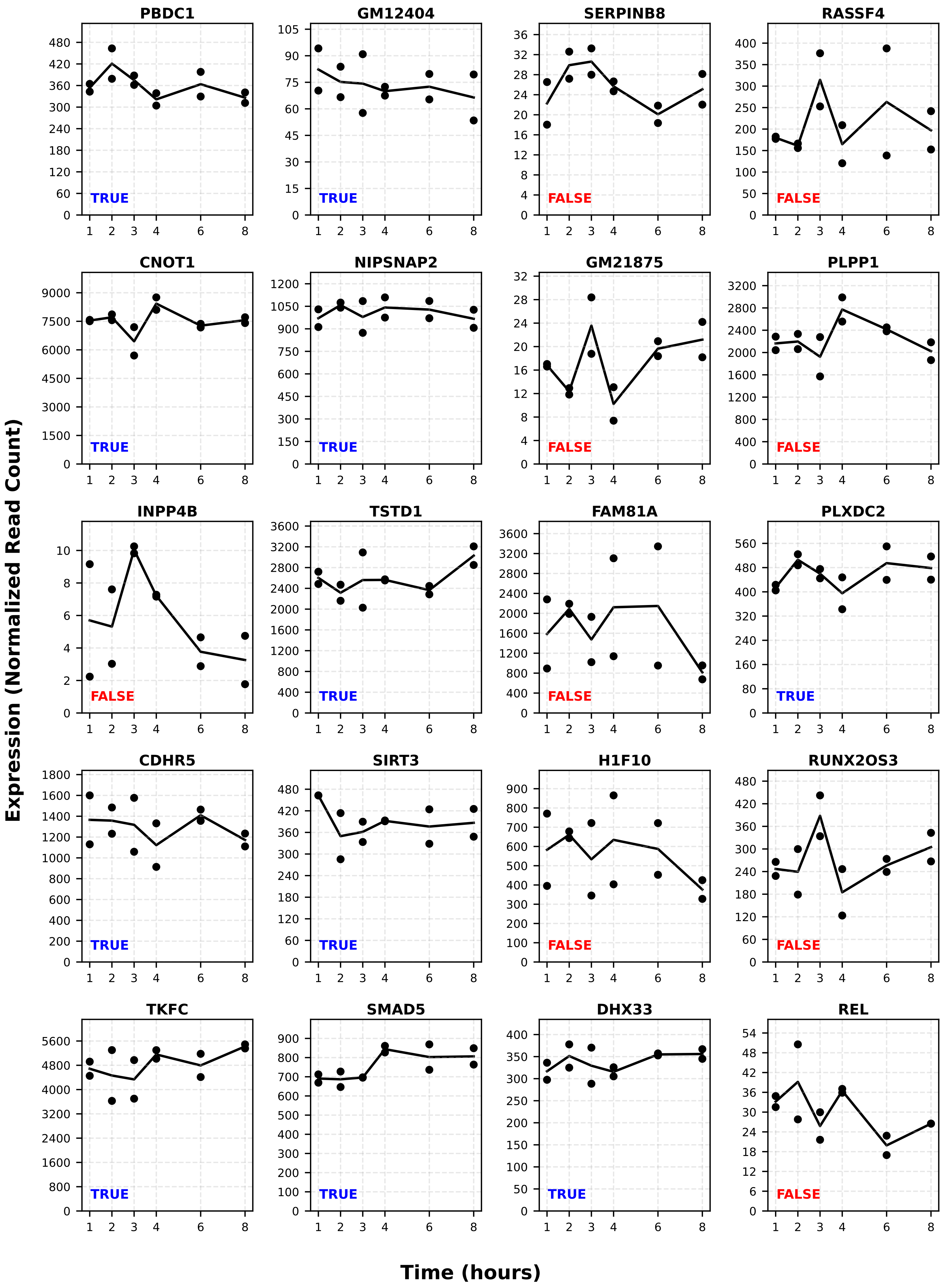

PBDC1
TRUE
GM12404
TRUE
SERPINB8
FALSE
RASSF4
FALSE
CNOT1
TRUE
NIPSNAP2
TRUE
GM21875
FALSE
PLPP1
FALSE
INPP4B
FALSE
TSTD1
TRUE
FAM81A
FALSE
PLXDC2
TRUE
CDHR5
TRUE
SIRT3
TRUE
H1F10
FALSE
RUNX2OS3
FALSE
TKFC
TRUE
SMAD5
TRUE
DHX33
TRUE
REL
FALSE
Expression (Normalized Read Count)
Time (hours)

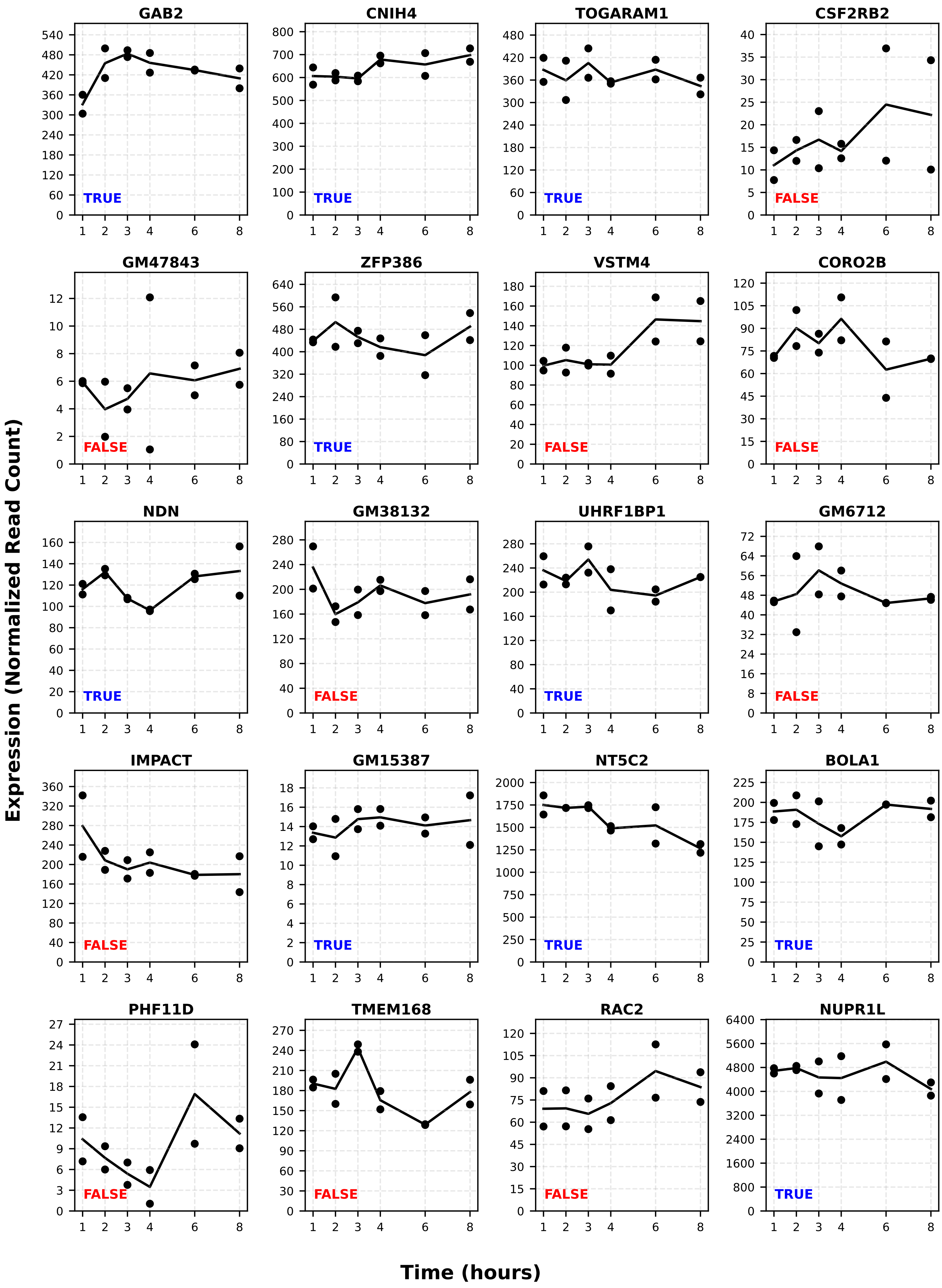
GAB2
TRUE
CNIH4
TRUE
TOGARAM1
TRUE
CSF2RB2
FALSE
GM47843
FALSE
ZFP386
TRUE
VSTM4
FALSE
CORO2B
FALSE
NDN
TRUE
GM38132
FALSE
UHRF1BP1
TRUE
GM6712
FALSE
IMPACT
FALSE
GM15387
TRUE
NT5C2
TRUE
BOLA1
TRUE
PHF11D
FALSE
TMEM168
FALSE
RAC2
FALSE
NUPR1L
TRUE
Expression (Normalized Read Count)
Time (hours)

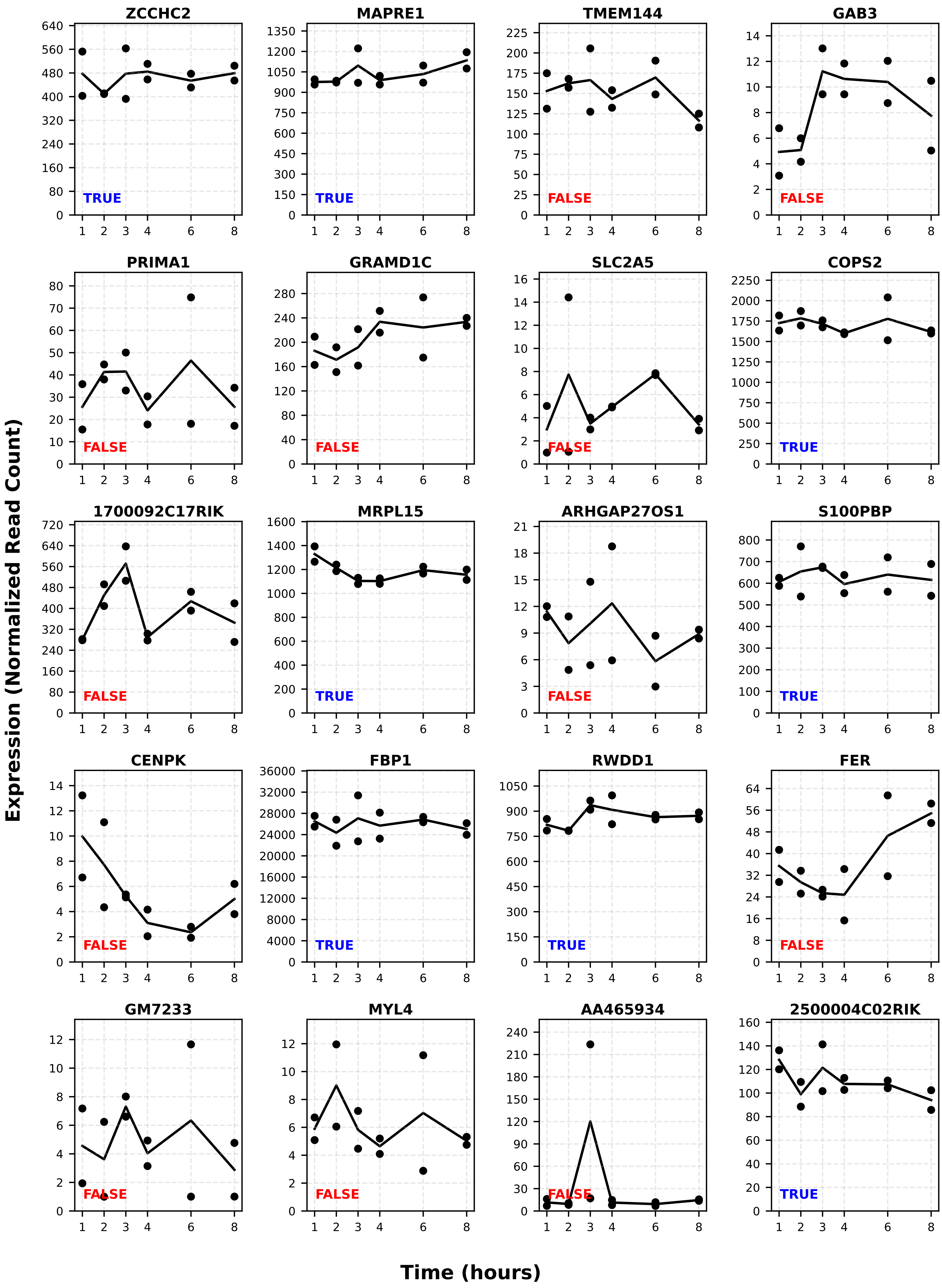
ZCCHC2
TRUE
MAPRE1
TRUE
TMEM144
FALSE
GAB3
FALSE
PRIMA1
FALSE
GRAMD1C
FALSE
SLC2A5
FALSE
COPS2
TRUE
1700092C17RIK
FALSE
MRPL15
TRUE
ARHGAP27OS1
FALSE
S100PBP
TRUE
CENPK
FALSE
FBP1
TRUE
RWDD1
TRUE
FER
FALSE
GM7233
FALSE
MYL4
FALSE
AA465934
FALSE
2500004C02RIK
TRUE
Expression (Normalized Read Count)
Time (hours)

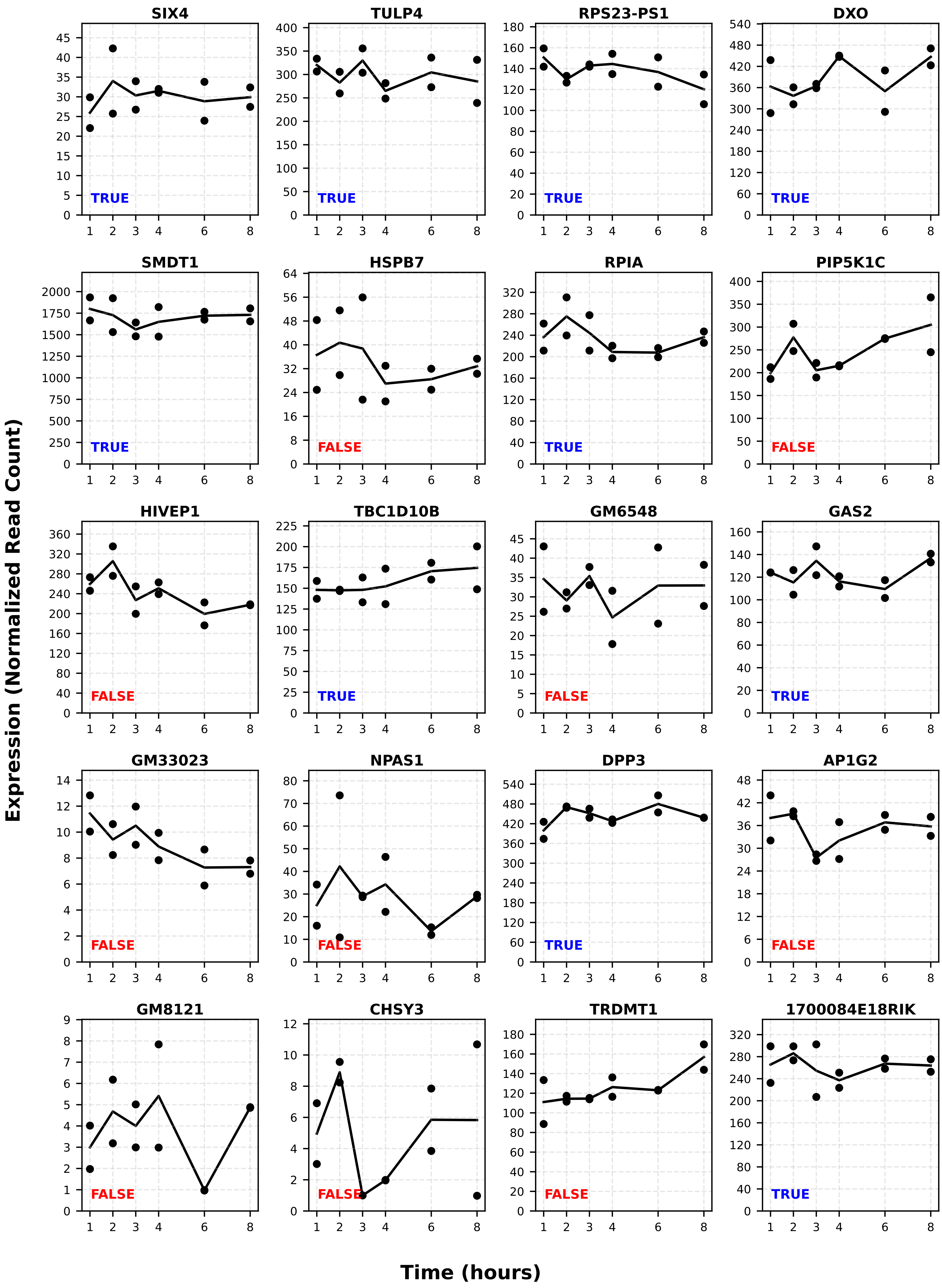
SIX4
TULP4
RPS23-PS1
DXO
SMDT1
HSPB7
RPIA
PIP5K1C
HIVEP1
TBC1D10B
GM6548
GAS2
GM33023
NPAS1
DPP3
AP1G2
GM8121
CHSY3
TRDMT1
1700084E18RIK
TRUE
FALSE
Expression (Normalized Read Count)
Time (hours)

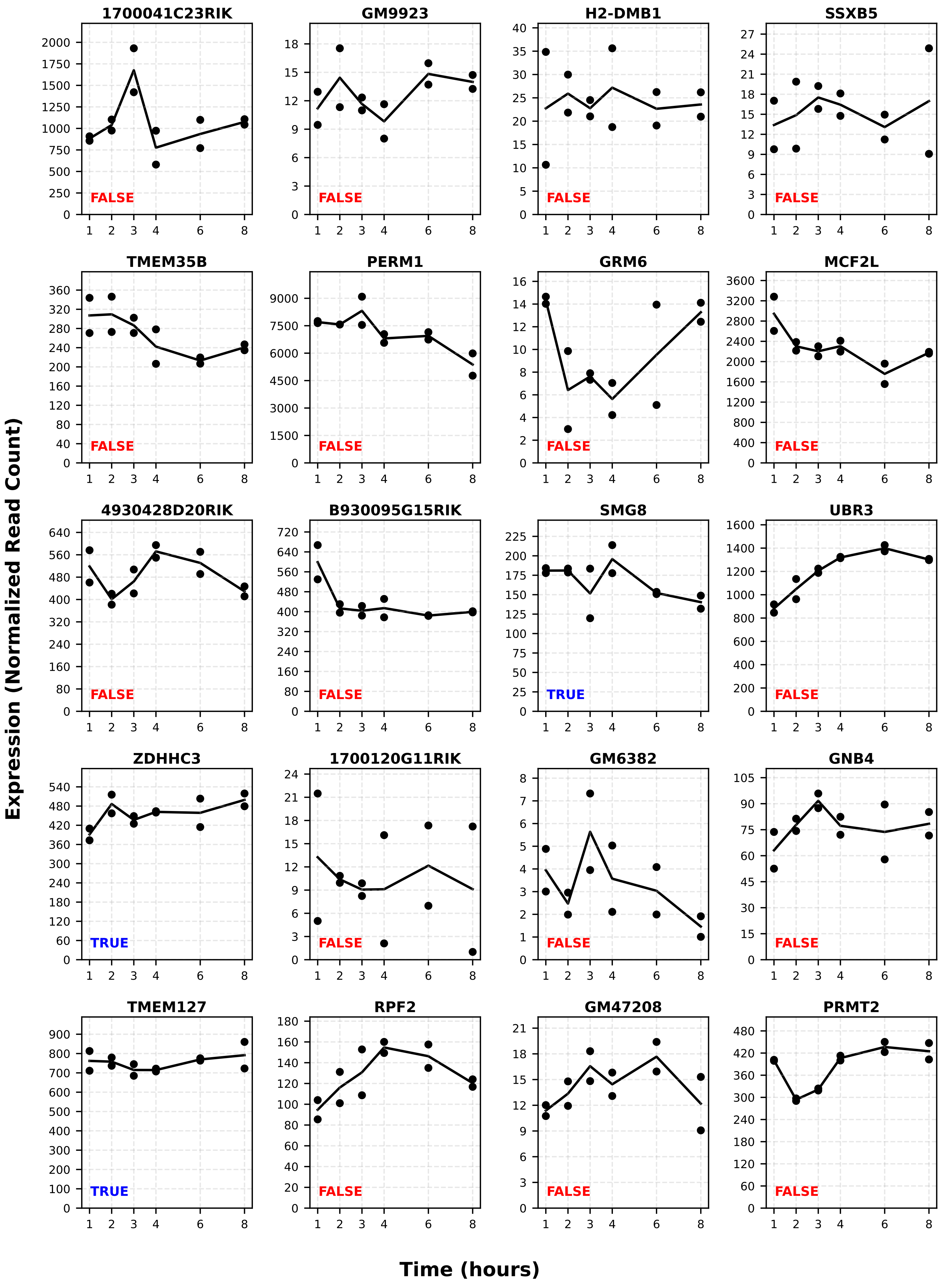
1700041C23RIK
GM9923
H2-DMB1
SSXB5
TMEM35B
PERM1
GRM6
MCF2L
4930428D20RIK
B930095G15RIK
SMG8
UBR3
ZDHHC3
1700120G11RIK
GM6382
GNB4
TMEM127
RPF2
GM47208
PRMT2
FALSE
TRUE
Expression (Normalized Read Count)
Time (hours)

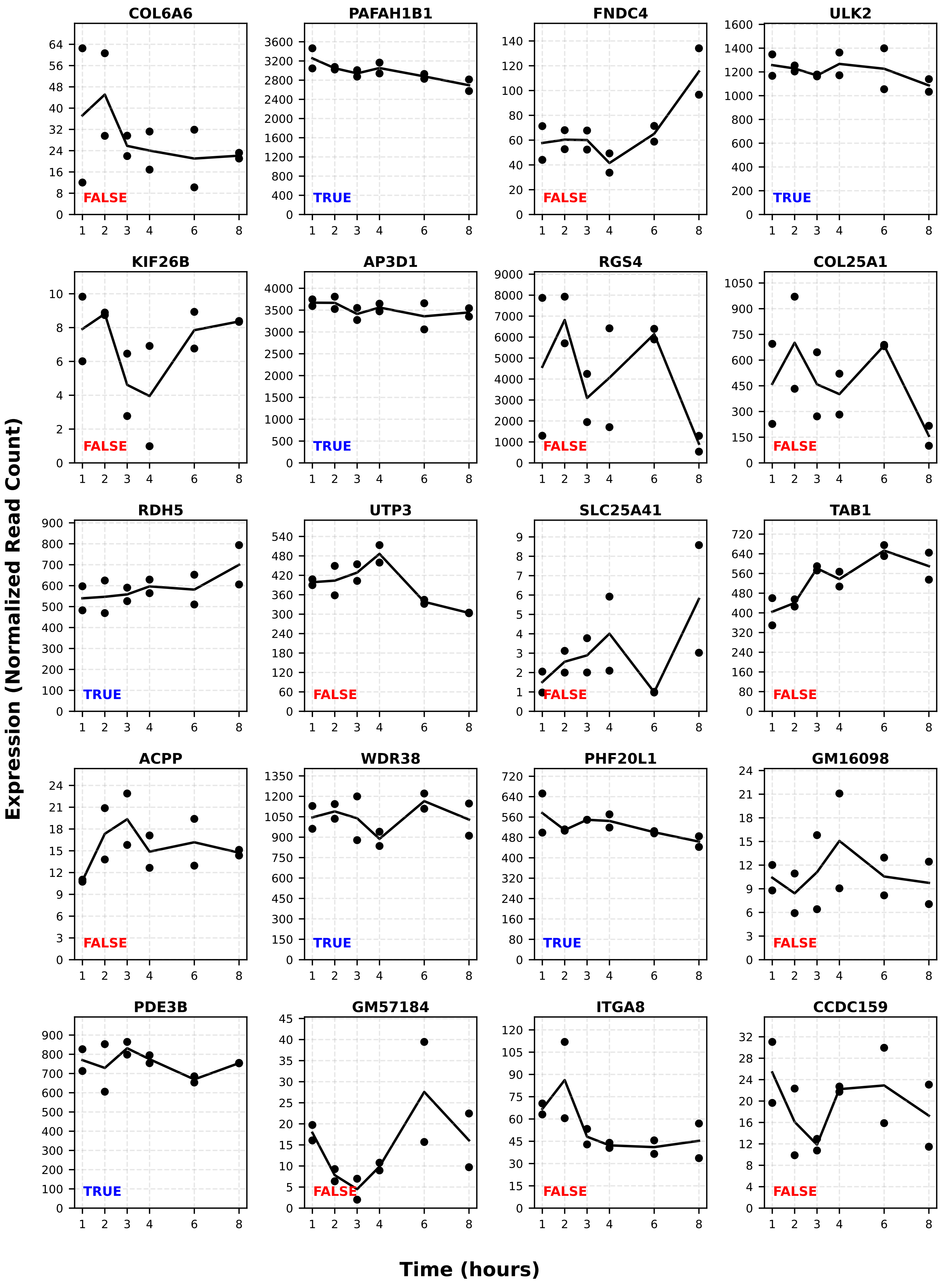
COL6A6
FALSE
PAFAH1B1
TRUE
FNDC4
FALSE
ULK2
TRUE
KIF26B
FALSE
AP3D1
TRUE
RGS4
FALSE
COL25A1
FALSE
RDH5
TRUE
UTP3
FALSE
SLC25A41
FALSE
TAB1
FALSE
ACPP
FALSE
WDR38
TRUE
PHF20L1
TRUE
GM16098
FALSE
PDE3B
TRUE
GM57184
FALSE
ITGA8
FALSE
CCDC159
FALSE
Expression (Normalized Read Count)
Time (hours)

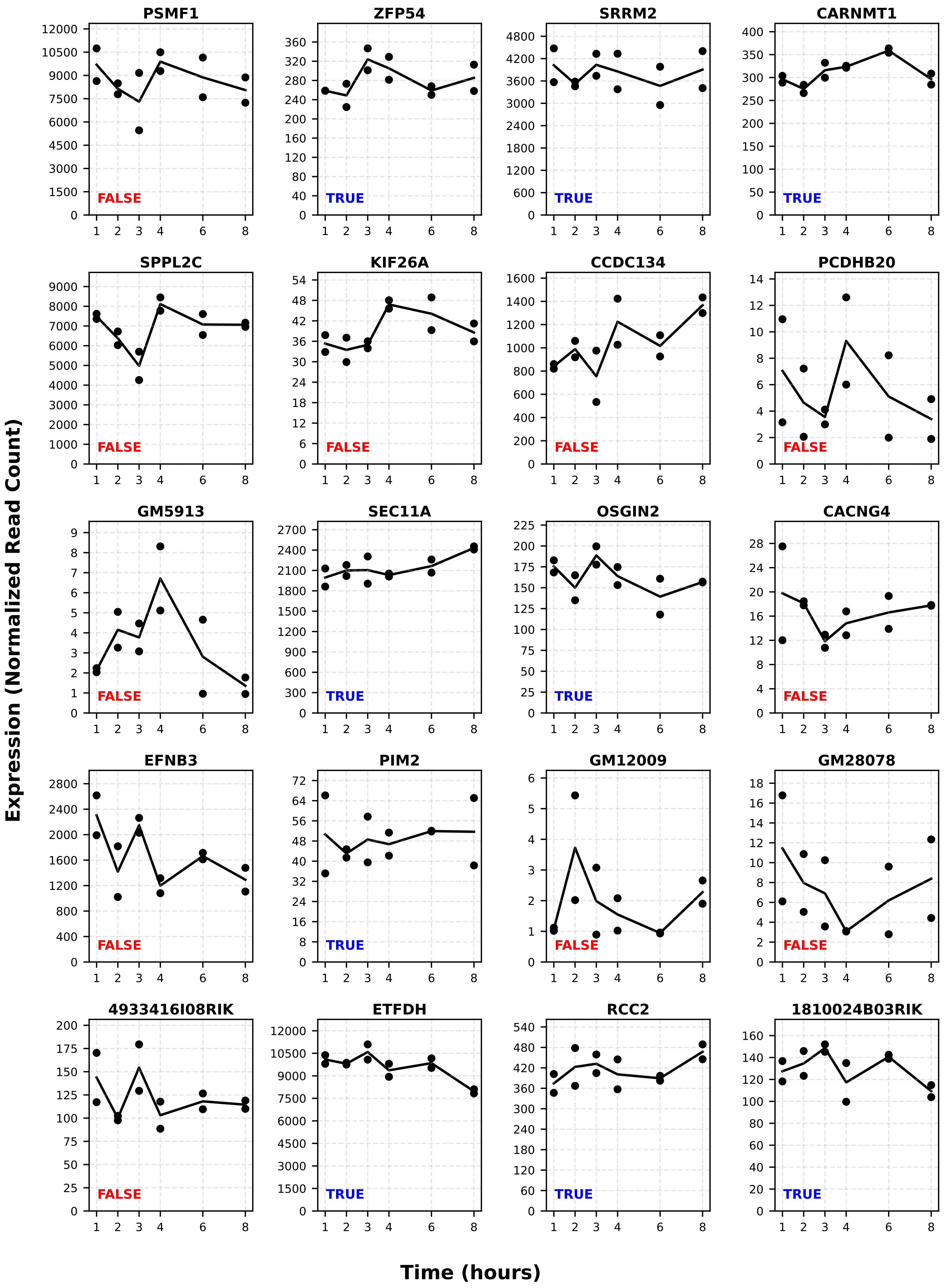
PSMF1
FALSE
ZFP54
TRUE
SRRM2
TRUE
CARNMT1
TRUE
SPPL2C
FALSE
KIF26A
FALSE
CCDC134
FALSE
PCDHB20
FALSE
GM5913
FALSE
SEC11A
TRUE
OSGIN2
TRUE
CACNG4
FALSE
EFNB3
FALSE
PIM2
TRUE
GM12009
FALSE
GM28078
FALSE
4933416I08RIK
FALSE
ETFDH
TRUE
RCC2
TRUE
1810024B03RIK
TRUE
Expression (Normalized Read Count)
Time (hours)

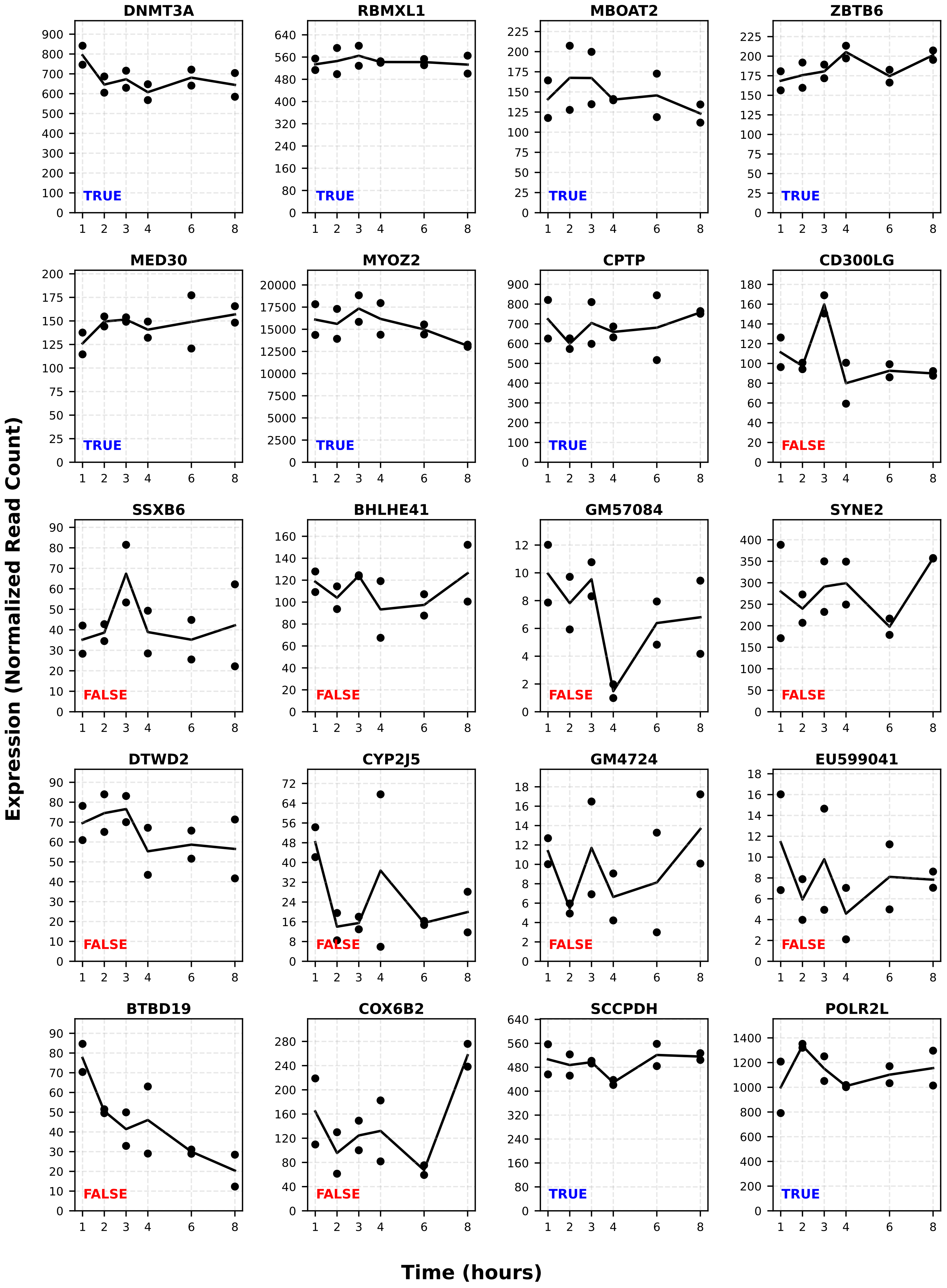
DNMT3A
RBMXL1
MBOAT2
ZBTB6
MED30
MYOZ2
CPTP
CD300LG
SSXB6
BHLHE41
GM57084
SYNE2
DTWD2
CYP2J5
GM4724
EU599041
BTBD19
COX6B2
SCCPDH
POLR2L
TRUE
FALSE
Expression (Normalized Read Count)
Time (hours)

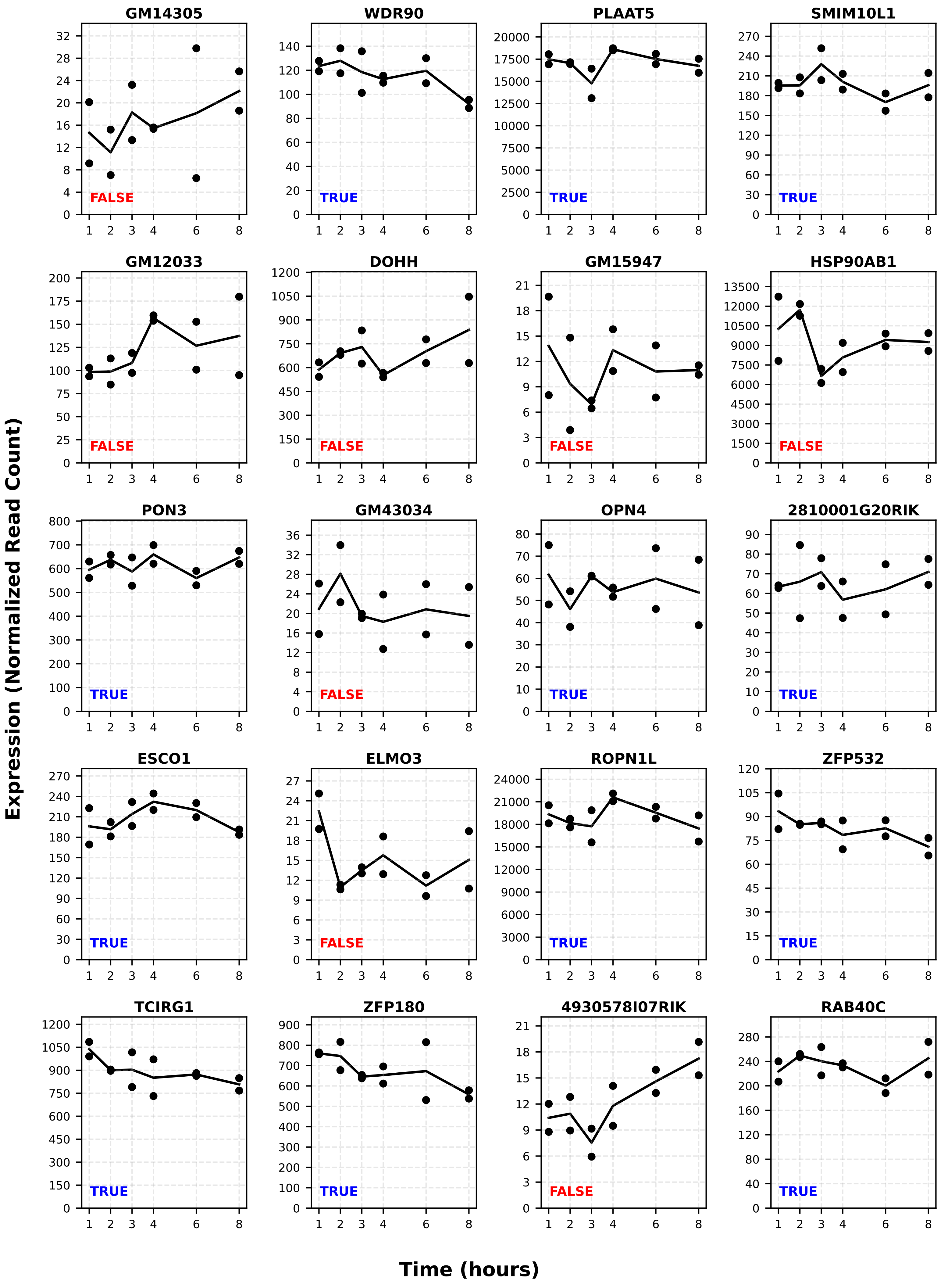
GM14305
FALSE
WDR90
TRUE
PLAAT5
TRUE
SMIM10L1
TRUE
GM12033
FALSE
DOHH
FALSE
GM15947
FALSE
HSP90AB1
FALSE
PON3
TRUE
GM43034
FALSE
OPN4
TRUE
2810001G20RIK
TRUE
ESCO1
TRUE
ELMO3
FALSE
ROPN1L
TRUE
ZFP532
TRUE
TCIRG1
TRUE
ZFP180
TRUE
4930578I07RIK
FALSE
RAB40C
TRUE
Expression (Normalized Read Count)
Time (hours)

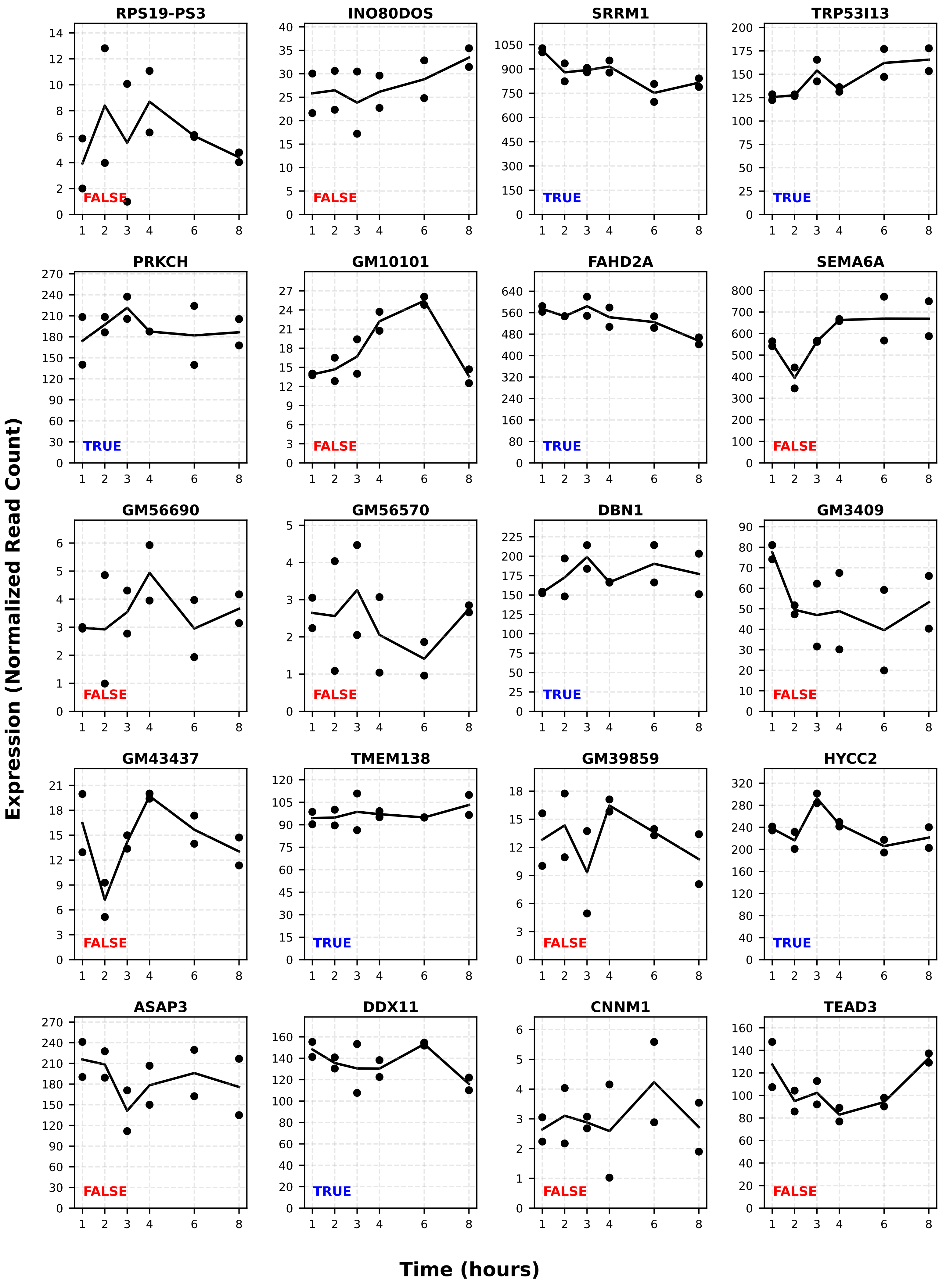

RPS19-PS3
FALSE
INO80DOS
FALSE
SRRM1
TRUE
TRP53I13
TRUE
PRKCH
TRUE
GM10101
FALSE
FAHD2A
TRUE
SEMA6A
FALSE
GM56690
FALSE
GM56570
FALSE
DBN1
TRUE
GM3409
FALSE
GM43437
FALSE
TMEM138
TRUE
GM39859
FALSE
HYCC2
TRUE
ASAP3
FALSE
DDX11
TRUE
CNNM1
FALSE
TEAD3
FALSE
Expression (Normalized Read Count)
Time (hours)

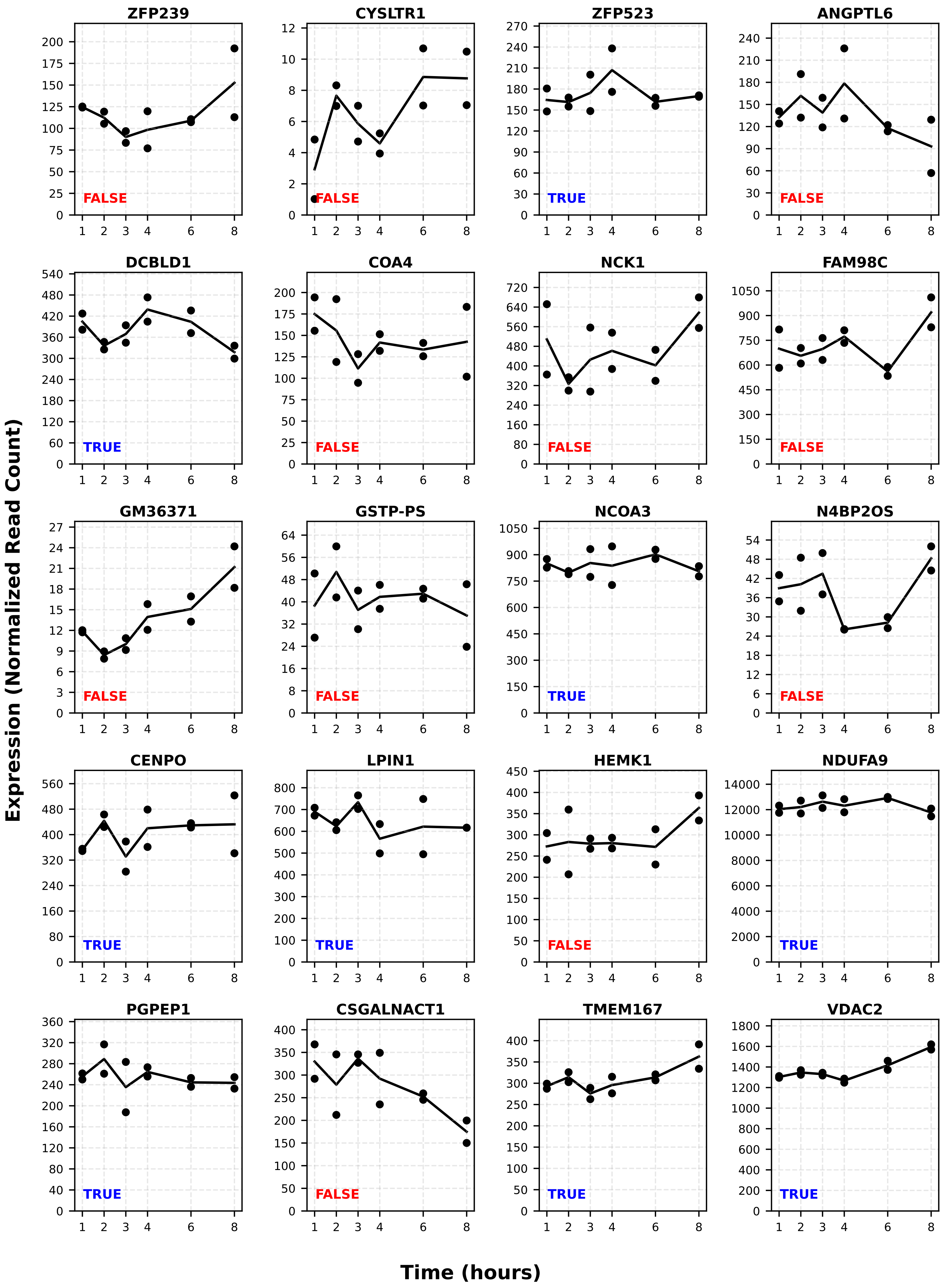

ZFP239
FALSE
CYSLTR1
FALSE
ZFP523
TRUE
ANGPTL6
FALSE
DCBLD1
TRUE
COA4
FALSE
NCK1
FALSE
FAM98C
FALSE
GM36371
FALSE
GSTP-PS
FALSE
NCOA3
TRUE
N4BP2OS
FALSE
CENPO
TRUE
LPIN1
TRUE
HEMK1
FALSE
NDUFA9
TRUE
PGPEP1
TRUE
CSGALNACT1
FALSE
TMEM167
TRUE
VDAC2
TRUE
Expression (Normalized Read Count)
Time (hours)

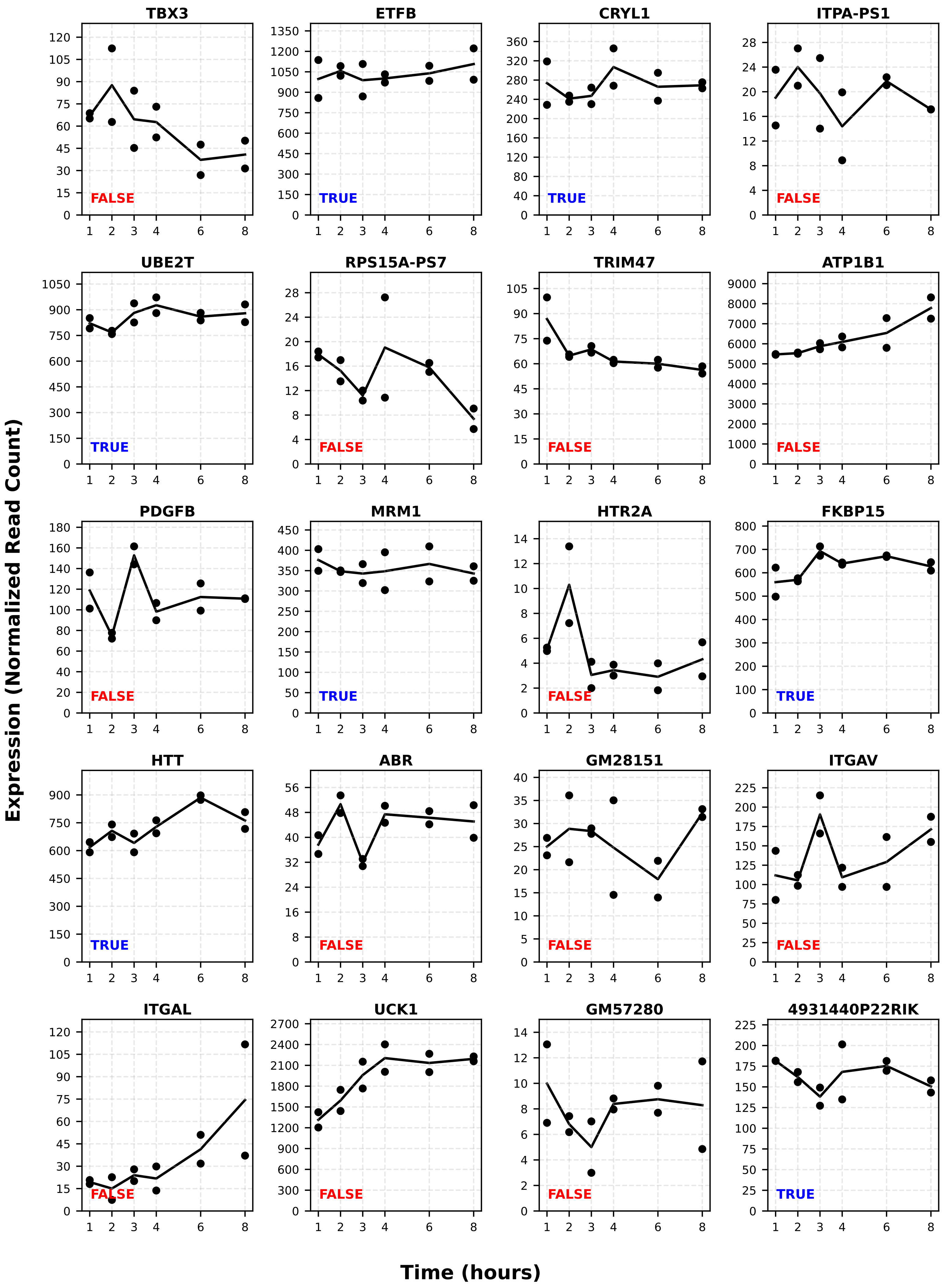
TBX3
ETFB
CRYL1
ITPA-PS1
UBE2T
RPS15A-PS7
TRIM47
ATP1B1
PDGFB
MRM1
HTR2A
FKBP15
HTT
ABR
GM28151
ITGAV
ITGAL
UCK1
GM57280
4931440P22RIK
FALSE
TRUE
Expression (Normalized Read Count)
Time (hours)

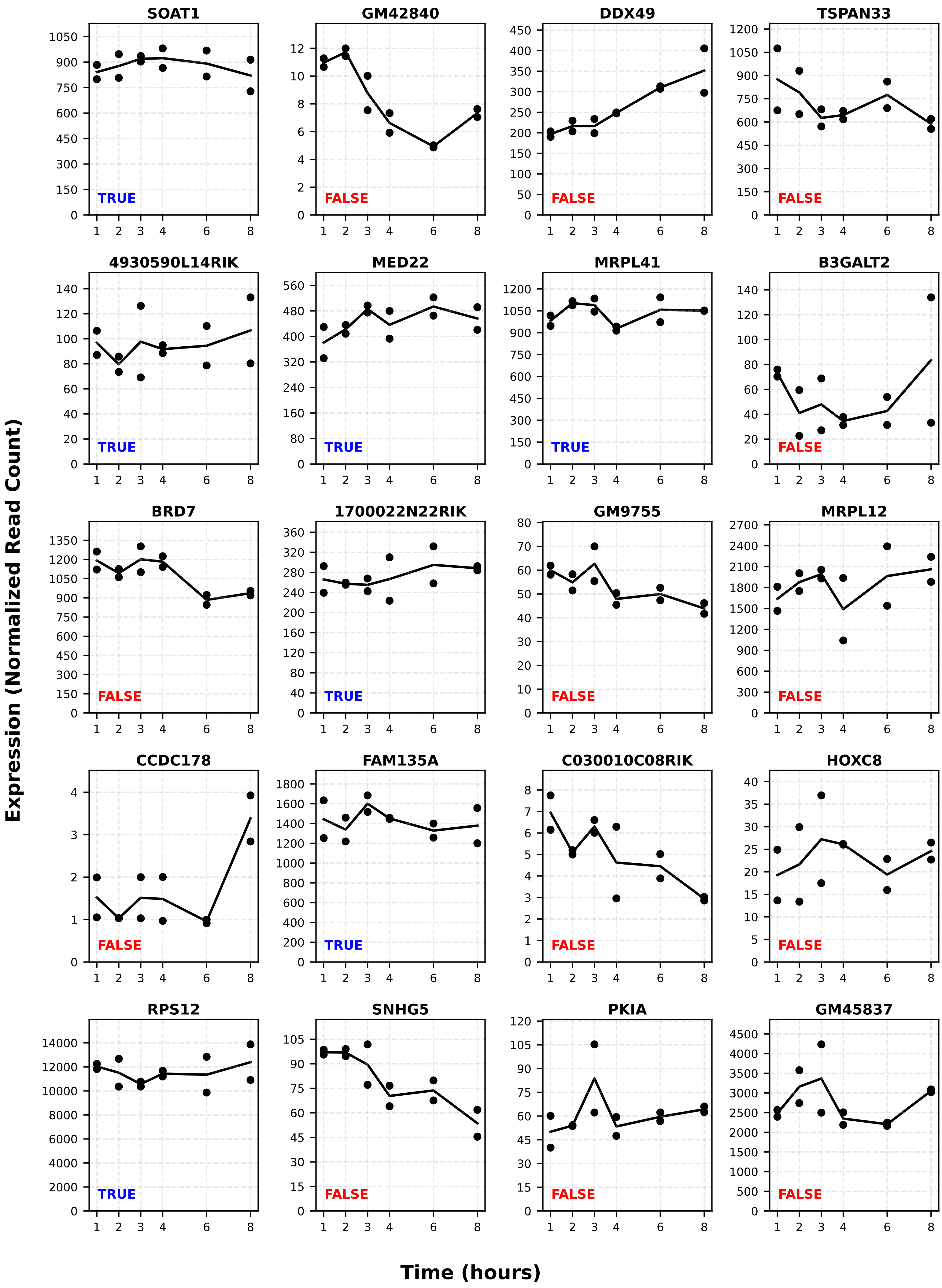
SOAT1
TRUE
GM42840
FALSE
DDX49
FALSE
TSPAN33
FALSE
4930590L14RIK
TRUE
MED22
TRUE
MRPL41
TRUE
B3GALT2
FALSE
BRD7
FALSE
1700022N22RIK
TRUE
GM9755
FALSE
MRPL12
FALSE
CCDC178
FALSE
FAM135A
TRUE
C030010C08RIK
FALSE
HOXC8
FALSE
RPS12
TRUE
SNHG5
FALSE
PKIA
FALSE
GM45837
FALSE
Expression (Normalized Read Count)
Time (hours)
