## Supplemental Figure 2 for "RNA degradation modulates unique aging-related gene expression in naked mole-rats"

### ***Example of RNA expression***

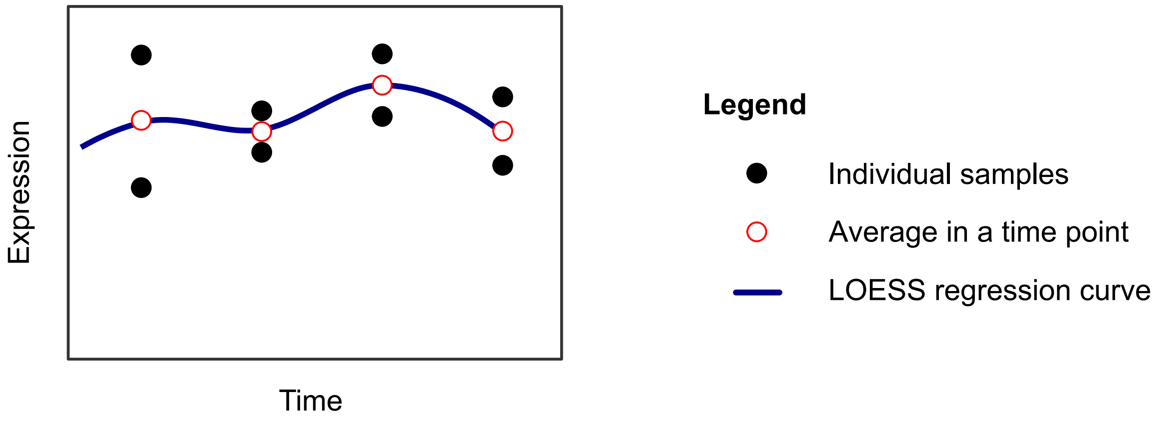


### ***The 3-layer test***

*i) Differential expression analysis*

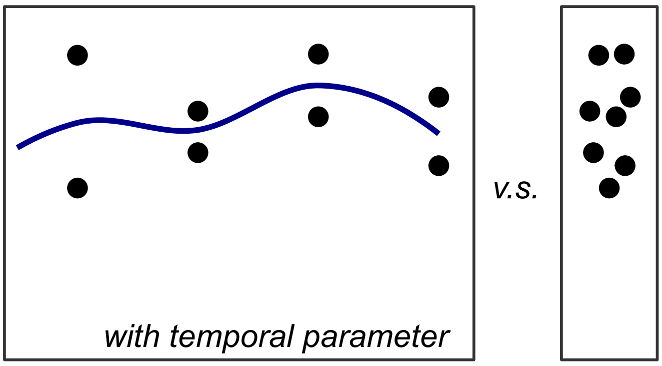


(Adjusted p-value > threshold$^{i}$ ) $\Longrightarrow$ (steady-state)

*ii) maximum of replicate differences*

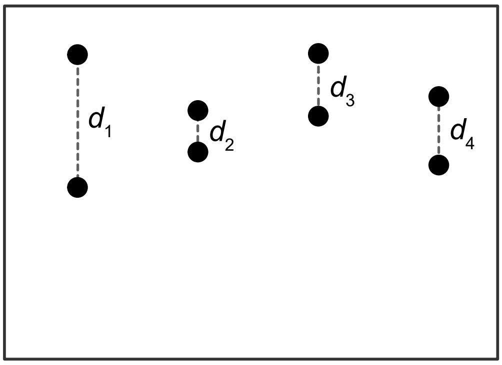


$$\left( \max_{k=1,\ldots,n} \{d_k\} < \text{threshold}^{ii} \right) \Longrightarrow \text{(steady-state)}$$

*iii) maximum of fold changes*

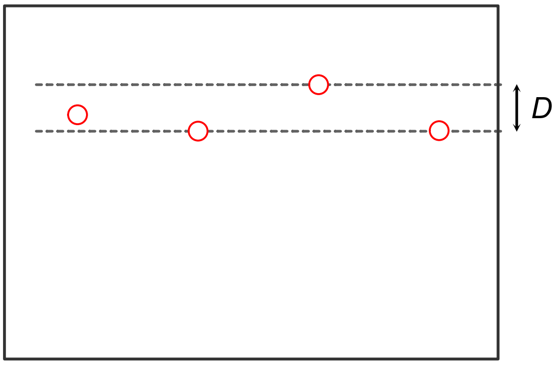


($D$ < threshold$^{iii}$ ) $\Longrightarrow$ (steady-state)
